# 3D confinement reshapes RNA folding and enhances circularisation in the Zika virus

**DOI:** 10.64898/2026.01.10.698690

**Authors:** J. K. Novev, J. Y. Lau, D. Marenduzzo, G. Kudla

## Abstract

Many RNA molecules function within confined environments, but the effect of confinement on RNA folding remains poorly understood. Proximity ligation experiments reveal altered long-range contacts in confined versus unconfined states, yet they do not explain how spatial constraints give rise to these differences. Here, we develop a physical modeling approach that incorporates proximity ligation data into coarse-grained molecular dynamics simulations to reconstruct RNA 3D structures under confinement. We test our model on the ∼ 11 kb genome of the Zika virus, comparing the folding in virions (confined) and in cells (unconfined). We observe that the probability of contact between two regions of the genome vs. linear distance follows different scaling laws in the confined and unconfined cases, in agreement with proximity ligation experiments. We find that genome circularization – an interaction that regulates replication in Zika – occurs more frequently under confinement in both experiments and simulations. Our model reveals that the formation of long double-stranded stems through stacking confers local nematic liquid crystalline order to the RNA, and predicts pseudoknot topologies consistent with those seen in crystallographic structures of shorter RNAs. These results provide insight into the mechanism through which confinement alters the ensemble of 3D structures accessible to a long RNA molecule.

**Significance statement:** Squeezing a polymer into a tight ball changes how it behaves: distant segments are forced into close contact, making ends more likely to meet. Similar principles apply to biopolymers, such as RNA, when confined within virus particles or other enclosures smaller than the polymer contour length, but there is no framework to model the effects of confinement on RNA folding. Here we develop such a framework, combining molecular dynamics simulations with experimental measurements of contacts within unconfined and confined RNA molecules. We find that RNA is glassy and exhibits liquid crystalline order, and that genome circularization, a long-range interaction between the two ends of the viral RNA important for virus replication, is facilitated under confinement.

---

RNA molecules often reside inside tightly confined spaces. This is true of viral genomes enclosed within protein capsids [1], of mRNAs and lncRNAs inside phase-separated RNP granules [2], and of RNA therapeutics encapsulated within lipid nanoparticles [3]. Despite the ubiquity of such systems, there is currently no general framework for predicting how confinement affects RNA folding.

Emerging experimental and theoretical evidence demonstrates that confinement can alter both the structure and the function of an RNA. The frequency and patterns of long-range interactions probed with proximity ligation assays differ between RNA virus genomes probed in cells and in virions – including Zika [1, 4], SARS-CoV-2 [5], and HIV [6]. A recent molecular dynamics study predicted new long-range contacts and pseudoknots upon encapsidation of the cowpea chlorotic mottle virus genome [7]. There are also notable differences in folding found by experiments on the lncRNA NORAD localized to the cytoplasm versus inside stress granules [8]. It has further been shown that modifying spatial confinement can affect RNA activity: for example, the catalytic efficiency of ribozymes is altered by encapsulation in lipid vesicles [9, 10]. From a thermodynamic perspective, confinement increases excluded-volume interactions and penalizes unfolded states, thereby perturbing the RNA energy landscape and favoring compact conformations [10, 11]. Although RNA structures are often modeled computationally using physical [12–14] and machine learning approaches [15, 16], these methods do not include confinement as an adjustable parameter. As a result, we currently lack tools for studying the effect of confinement on RNA folding.

We address this gap with a data-driven polymer modeling framework that combines base-pairing contact frequencies from proximity ligation experiments [1, 4] with coarse-grained molecular dynamics to construct 3D models of RNA folding. This approach enables us to simulate the folding of large RNAs and compare structural ensembles with and without confinement. We focus on the genome of the Zika virus, a ∼ 11 kb single-stranded RNA that has been extensively profiled using proximity ligation both in cells (unconfined conditions) [1] and inside virions (confined conditions) [4]. The Zika genome contains functionally important RNA secondary structure elements, including circularization motifs that regulate the switching between translation and replication [17], as well as pseudoknots that stabilize subgenomic RNAs [18, 19] and suppress RNAi-based innate immunity in the host during infection [20]. These features make Zika a biologically relevant model for studying how confinement shapes structure and function of RNA. Our results show that confinement reshapes long-range RNA contacts, promotes circularization, and alters pseudoknots and structural dynamics in the Zika virus, establishing a framework for modeling RNA folding under physical constraints.

The approach that we describe here has advantages over existing methods for studying long RNAs and viral genomes. First, machine-learning-based approaches [15, 16] are typically trained on known structures for particular RNA families, and their predictions do not generalize well to RNAs from other families [21] that are less well represented in the literature, such as viral RNAs. Second, although molecular dynamics simulations have previously been used to study viruses [22, 23], including Zika [24], most studies have focussed on the viral envelope and capsid rather than the viral genome. Third, experimental methods developed in the last several years have brought advances in resolving the structure of viral RNAs: e.g., cryo-electron microscopy (cryo-EM) has allowed the mapping of the genome of bacteriophages such as MS2 [25, 26] and AP205 [27]. However, cryo-EM reconstruction is more difficult for viruses such as Zika, where RNA-protein binding is less well-defined [28] and the genome is not asymmetrically anchored to the capsid.

## Results

### Experimental contact patterns and probability are fundamentally different under confinement

We use RNA proximity ligation datasets from the Zika virus to study the influence of physical confinement on RNA folding. In proximity ligation, RNA is crosslinked, fragmented, ligated, and sequenced to identify base-pairing contacts [29]. We use COMRADES [1], SPLASH [4], and PARIS [30] data from several strains of Zika, obtained under three conditions: in cells, in vitro refolded, and in virions. For consistency, we reanalyze all datasets using a unified pipeline, combining experimental replicates to generate intramolecular contact maps. When applied to nearly identical virus strains (*>* 99.5% sequence identity) under the same conditions, SPLASH, PARIS and COMRADES recover similar patterns of interactions (Suppl. Fig. S1), but COMRADES recovers more than 10 times as many in-cell reads as PARIS or SPLASH (Suppl. Figs. S1-S3), likely because COMRADES includes an enrichment step where Zika RNA fragments are pulled down using biotinylated oligonucleotides. We consider in-cell and in-vitro-refolded RNA as unconfined and in-virion as physically confined because it is only in the virion, but not in cells or in vitro, that the RNA is constrained to an enclosure of size smaller than the RNA’s contour length.

Across COMRADES, PARIS, and SPLASH, maps from cytoplasmic and refolded RNA are dominated by local interactions, visible as strong signal close to the diagonal (Suppl. Figs. S4-S6). In contrast, we find that virion-derived RNAs (for which only SPLASH data is available) exhibit numerous distinct long-range contacts (Fig. 1). The enrichment of long-range interactions in virion RNA can be captured by the probability of contact, *p*_contact_, which measures the probability that in an ensemble of configurations two segments of a polymer chain with linear separation *l* are in contact (note this means base-paired in experiments). In polymer physics this probability often follows a power law,

**Figure 1.**
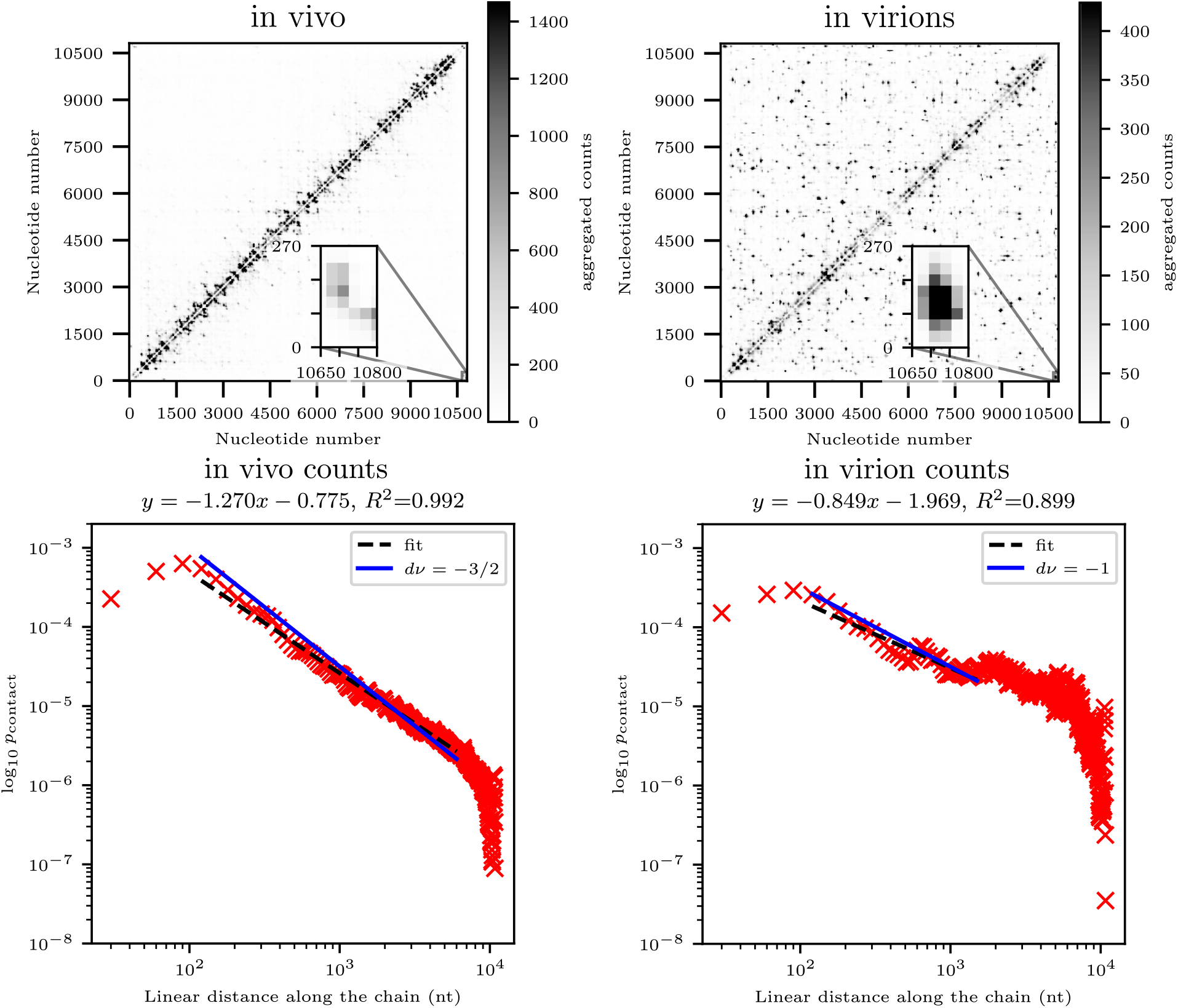
The contact patterns (top row) and curves of contact probability vs. genomic distance (bottom row) are strikingly different for data recorded in cells (left column) and in virions (right column). Contact maps are symmetrized to show aggregated counts corresponding to signal from chimeric reads of both orientations (see Methods: Correlations between contact maps) and coarse-grained into bins of length 30 nucleotides. Contacts with aggregated counts equal to or above the 99th percentile for the specific dataset are shown in black. Data for in vivo conditions from [1] and for virions from [4]. The probability of contacts between segments as a function of their separation for the in vivo data (left) is close to the limiting law for an ideal chain (slope −3*/*2), whereas for the data collected under in virion conditions (right) the exponent is approximately equal to −1. Note the non-monotonic dependence of *p*_contact_ on distance - this is due partly to limitations of detecting short-range contacts in proximity ligation methods (see Methods: Contact probability exponents) and partly to base-pairing between the two ends of the viral genome (see Results: Simulations and experiments demonstrate higher circularization frequency under confinement).

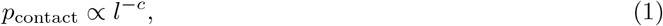

where *c* is a contact exponent which depends on universal features of the underlying polymer model rather than molecular detail. The contact exponent is linked to the spatial dimension *d* and the metric exponent *ν* [31] via the scaling law *c* = *dν*, as a result of which the value of *c* can be predicted for a number of relevant models in *d* = 3. Thus, for ideal chains, i.e., polymers without self-interactions, 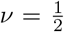 and 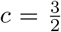 [32]; for self-avoiding chains, 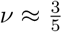 and 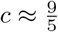 [31]^1^, while *c* = 1 for a fractal (crumpled) globule, which is observed when a polymer chain collapses quickly, or in the presence of topological constraints that disallow the passing of regions of the chain across each other [32].

Remarkably, as seen in Figure 1, our analysis of the experimental data yields slopes (−*c*) of −1.27 and −0.85 at intermediate distances for the unconfined (in vivo) and confined (in virion) states respectively. The two values are significantly different, and we note they are numerically close to the theoretical values for an ideal chain (−3*/*2) and a fractal globule (−1) respectively. Data obtained under the same conditions via different proximity ligation methods (COMRADES [1], SPLASH [4] and PARIS [30]) yields similar values of *c*, see Suppl. Fig. S4, indicating that confinement, rather than experimental differences, drives the observed variation in *c*. This suggests that confinement in the viral capsid sharply changes the nature of the folding of RNA. The fact that RNA in the unconfined cellular environment has a contact exponent compatible with that of a random walk is at first surprising; however, a recent numerical analysis [34] has found that the approximate metric exponent of RNA is compatible with 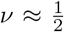, as in a random walk. We henceforth use data from COMRADES from the Brazil strain as representative of the unconfined state because it has the highest coverage for these conditions, and SPLASH data from the nearly identical (*>* 99.7% sequence identity) French Polynesian strain for the confined state.

Next, we test whether current RNA secondary structure prediction algorithms could recapitulate these results. We generate Boltzmann ensemble base-pairing probabilities using RNAfold (ViennaRNA) [12], LinearPartition [35] and UNAFold [36]. Across all predictions, the fitted scaling exponent varies in the range *c* ∼ 1.2 − 2.2 for both the Zika sequences used for in vivo experiments (KX197192.1) and ones in virions (KJ776791.2), with typical values close to 1.5, similar to what is observed for unconfined RNA see Suppl. Table S1). Incorporating input from NAI-MaP assays [4], which provide information about the reactivity of nucleotides in the viral genome, into RNAfold and LinearPartition predictions, results in values of *c* that are closer to those determined from in vivo experiments rather than to those performed in virions, see Suppl. Table S1 and Suppl. Fig. S7 for details. These results highlight that spatial confinement in the virion dramatically reshapes the folding of the viral genome and requires a different modelling approach to the ones in current RNA structure prediction algorithms. We propose one such approach in this study.

### A data-driven polymer model for RNA folding reproduces the effect of confinement on contact patterns

To determine the effect of spatial confinement on the patterns of long-range contacts in viral RNA folding, we developed a coarse-grained molecular dynamics model of RNA folding with and without confinement (Fig. 2), implemented in LAMMPS [38]. The model, described in detail in the Methods, represents the RNA as a bead-spring polymer with one bead per nucleotide. Base pairs form and break dynamically based on proximity and a contact probability table derived from structural ensemble predictions or experimental data [Fig. 2(a)]. Paired regions have higher bending stiffness that reflects the increased rigidity of double-stranded RNA [39, 40] [Fig. 2(b-c)]. Confinement within the Zika virion is modeled by a potential that restricts the polymer to a volume that corresponds to that of the inner capsid [Fig. 2(d)]. As we discuss in more detail below, the final core ingredient of the model is a simplified account of base-pair stacking [Fig. 2(b)], which promotes the formation of long stems. We explore different versions of the model varying in whether stacking is included and in the input used for base-pairing probabilities. For the main version of the model - 3D COMRADES - we estimate the latter from experimental data, whereas in 3D Vienna, we use ensemble base-pairing probability predictions from RNAfold [12]; we provide details of the implementation in the Methods.

**Figure 2.**
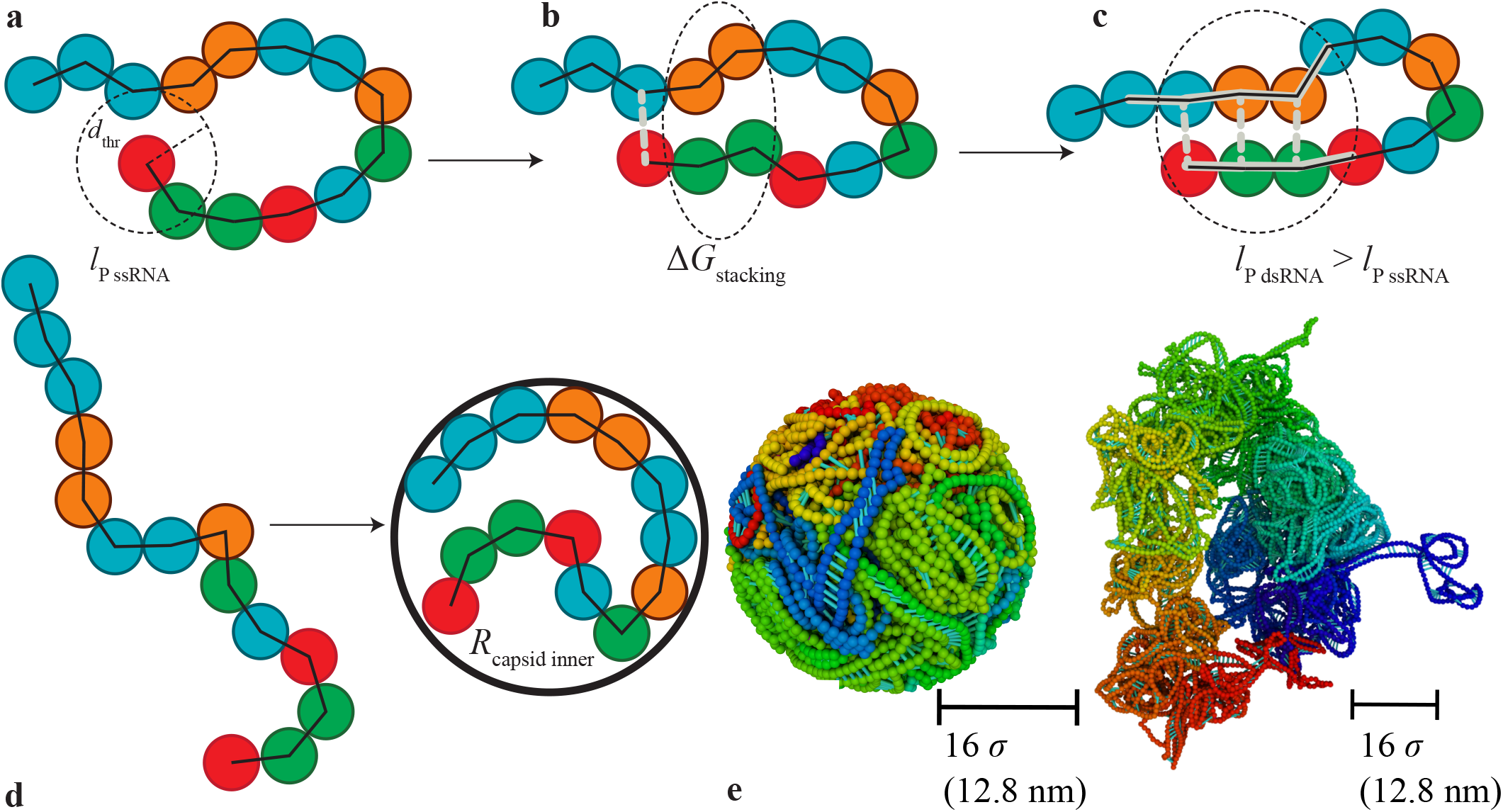
Cartoon of the main features of the model. Coarse-grained beads, each of them representing a ribonucleotide, are connected by permanent bonds (black lines) and angle bonds that confer bending rigidity to the RNA chain. If beads that can form a pair come within a distance *d*_thr_ of each other, the algorithm stochastically introduces a harmonic bond representing a base pair between them (**a**). Base-pair formation is more likely next to existing base-pairs due to stacking (**b**). Angle bonds between paired bases have a higher bending rigidity (stiffened angle bonds highlighted in gray), reflecting the higher stiffness of double-stranded RNA (**c**). The algorithm updates the list of base pairs at regular intervals, removing bases separated by a distance greater than *d*_thr_; pairs also break randomly with a probability of 1 - *w*_ij_, see Methods. Snapshots from simulations illustrate the spatial arrangement of the viral RNA with and without confinement (**e**); scale bar length approximately equal to the radius of the inner capsid of the Zika virus (**d**). 3D simulation snapshots here and in all other figures show beads approximately to scale and are rendered with the Open Visualization Tool (OVITO) [37]. Snapshots shown here do not show obvious tree-like structures, which are instead apparent in simulations of unconfined RNA based on Vienna base-pairing (SI, Fig. S28) because the data we show in the main text correspond to base pairing in spatially confined or *in vivo* situations..

**Figure 3.**
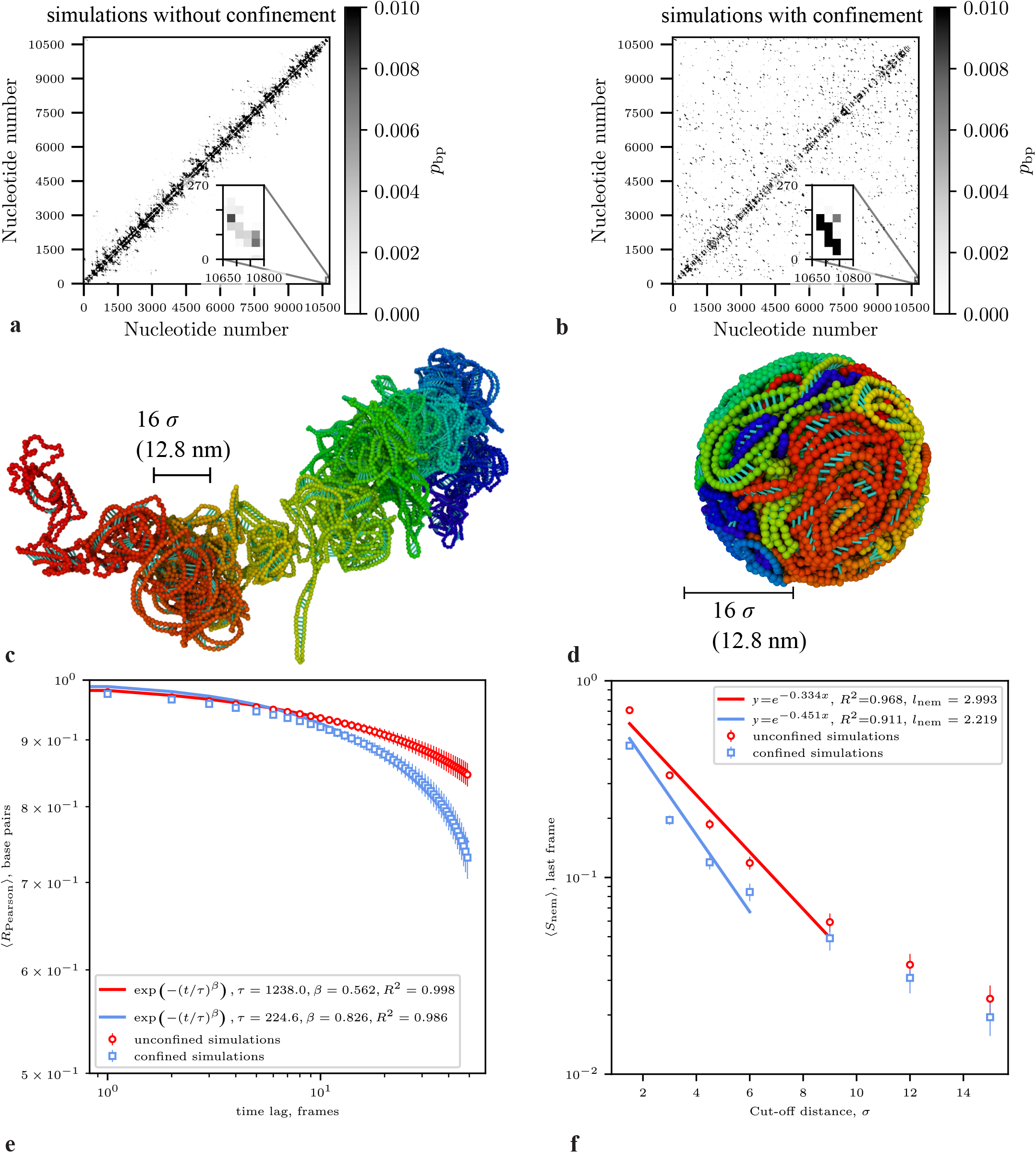
Simulations with stacking reproduce experimental contact maps, and predict glassy dynamics and nematic order in folded RNA. (**a,b**): Simulated contact maps with insets showing the circularization interactions for confined and unconfined RNA. (**c,d**): Representative snapshots for each case (nucleotides coloured blue to red according to their index). In addition to producing contact maps that correlate well with the experiment (Table 1), the simulations predict pseudoknots with a genus that is close to the expectation for an RNA with the length of the Zika genome [41] (Table 2). (**e**): Ensemble-averaged correlation coefficients between base-pairing contact maps for different frames. Persistent correlations between base-pairing contacts across frames in the simulations indicate glass-like behaviour both with and without confinement. The ensemble- and time-averaged ⟨*R*_Pearson_⟩ between the flattened matrices of base-pairing contacts vs. time lag calculated for ensembles of unconfined simulations (red) and confined ones (blue) follows a stretched exponential dependence, suggesting glassy behaviour, with a longer correlation time *t* for the unconfined case. ⟨*R*_Pearson_⟩ is calculated by averaging *R*_Pearson_ for all frames separated by a given time lag, then averaging over this set. (**f**): Nematic domains have a characteristic length of ∼ 3*s* for the unconfined simulations with stacking and COMRADES input, as demonstrated by the fit of average local nematic order in the last simulation frame vs. cut-off. Compare this result with the smaller length scale of nematic organization in confined simulations with stacking in the same plot. Scale bar length in simulation snapshots approximately equal to the radius of the inner capsid of the Zika virus. Ensembles of 100 simulations for all cases, coarse-graining with bin size of 30 nt (a-b) and 10 nt (c-d).

**Figure 4.**
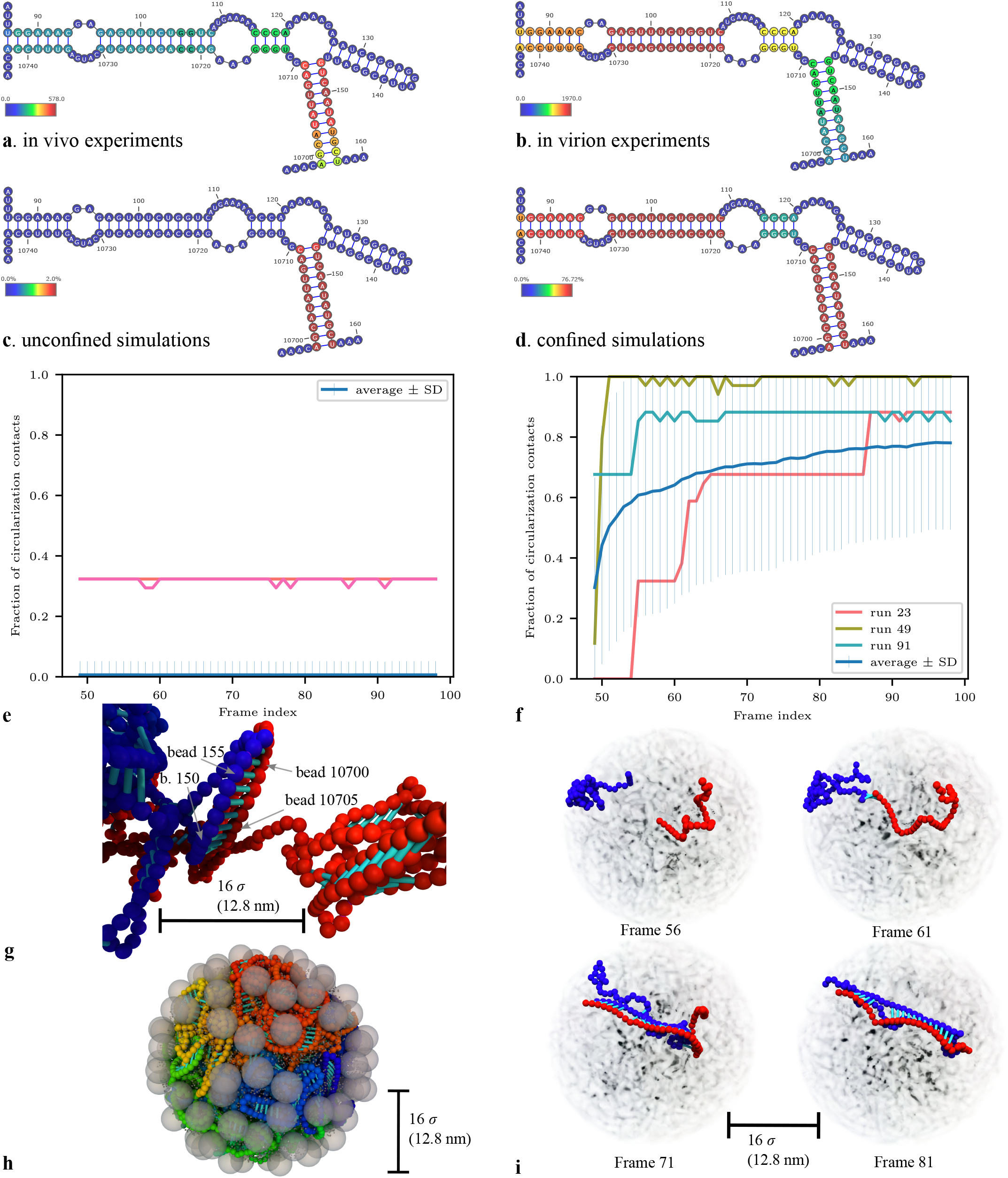
Circularization is enhanced by confinement and stem formation. (**a-b**): secondary structure diagrams of the circularization contacts specified in [17]; coloured by: left - proximity ligation *C*_ij_ scores calculated for in vivo conditions [1], right - *C*_ij_ scores for data from virions [4]. (**c-d**): colours based on ensemble- and time-averaged base-pairing probabilities from: left - unconfined simulations and right - confined simulations, both simulation types involving stacking and *C*_ij_ score input from the corresponding condition. The scores for the base pairs in the stem comprising the short-range interaction between nucleotides 127-135 and 146-138 are set to zero throughout. Third row: fraction of the contacts shown in the top panel that exist in an ensemble of unconfined (**e**) and confined (**f**) simulations. Lines with error-bars: ensemble averages; representative lines for individual simulations. **g**: snapshot from an unconfined simulation with unconfined experimental base-pairing *C*_ij_ score input and stacking with beads colored according to their index, going from blue (1) to red (10807). This snapshot displays one of the circularization stems that induce local nematic order. **h**: Snapshot from a simulation with electrostatics, an SBCG representation of the capsid (large grey beads) and sodium ions (small grey beads). **i**: snapshots from a confined simulation that illustrates the nucleation of a circularization stem and its dynamic evolution.

We evaluate our models by testing whether they generated plausible base-pairing patterns (based on stem length, fraction of paired bases, and absence of non-stem-like scattered long-range pairs); whether they recapitulate the scaling exponents observed experimentally; and the correlation of predicted base pairs with experimental data. The model that correlates best with experiments inputs a set of contact probability scores estimated from proximity ligation data, and does not involve stacking (Suppl. Fig. S8). The Pearson correlation coefficient is 0.709 for unconfined simulations (compared to experiments in vivo [1]), and 0.533 for confined simulations (compared to experiments in virion [4]); correlation coefficients are given in Table 1. Individual interactions such as those corresponding to circularization between the two ends of the viral genome (see inset of Suppl. Fig. S8) are also reproduced in simulations.

**Table 1:** *R*_Pearson_ measuring the correlation between different simulated contact maps and experimental chimeric count data. Left: unconfined simulations vs. in vivo data, right: confined simulations vs. in virion data. For simulations with base-pairing, *R*_Pearson_ measures the correlation of the simulated ensemble- and time-averaged base-pairing probabilities with the experimental counts. For simulations without base-pairing, *R*_Pearson_ measures the correlation between the matrix of ensemble- and time-averaged contacts defined to occur when the centres-of-mass of two bins are within *d*_cont_ of each other. Simulated and experimental maps are coarse-grained with a bin width of 30 nt.

| Type of prediction<br>for sequence<br>KX197192<br>(used for in vivo<br>experiments) | confinement | $R_{\text{Pearson}}$ vs<br>in vivo<br>counts | Type of prediction<br>for sequence<br>KJ776791.2<br>(used for in virion<br>experiments) | confinement | $R_{\text{Pearson}}$<br>vs in<br>virion<br>counts |
| --- | --- | --- | --- | --- | --- |
| no base-pairing,<br>contact at $d_{\text{cont}} = 10\sigma$ | — | 0.44 | no base-pairing,<br>contact at $d_{\text{cont}} = 10\sigma$ | + | 0.14 |
| RNAfold [12] prediction | — | 0.19 | RNAfold prediction | — | 0.14 |
| 3D Vienna | — | 0.12 | 3D Vienna | + | 0.10 |
| 3D COMRADES, no stacking | — | 0.71 | 3D COMRADES, no stacking | + | 0.53 |
| 3D COMRADES, stacking | — | 0.53 | 3D COMRADES, stacking | + | 0.34 |
| — | — | — | 3D COMRADES, stacking,<br>electrostatics, capsid | + | 0.37 |

Suppl. Fig. S8 illustrates contact map patterns that recapitulate the experimental ones, contact probability (with contacts defined through base pairing) vs. genomic distance curves that display a marked difference between the unconfined (*c* = 2.097) and the confined regimes (*c* = 0.910) in terms of contacts defined through interbead distances.

However, the high degree of correspondence between experiments and time- and ensemble averaged simulated contact maps in this case comes at the price of having unrealistic RNA secondary structures in the individual frames. In the absence of stacking, the stems within these simulations are predominantly of length 1-3 nt, frequently contain scattered long-range pairs, and many base-pairing interactions are highly stretched as seen in the snapshots in Suppl. Fig. S8, where base pairs are represented by cyan cylinders. This is further illustrated in Suppl. Figure S9, which shows the length of the harmonic bonds that we use to model base pairs vs. time.

### Stacking leads to physically realistic RNA structures with multiple pseudoknots

We address the issues with our initial model evident in Suppl. Fig. S8 by introducing an energy gain due to base-pair stacking [Fig. 2(c) and Methods], as well as an energy penalty for pairs connecting neighbouring bases to distant regions of the molecule.

As we demonstrate in the snapshot in Fig. and Suppl. Fig. S10, stacking removes the problems with bond strain and stem length distributions whilst retaining base-pairing patterns that recapitulate those seen in the experiment – albeit with decreased correlation coefficients (*R*_Pearson_ = 0.529 for the unconfined case, and 0.335 for the confined case, see Table 1). In terms of contact probability curves, these remain very different for the unconfined case and the confined one, with the latter again consistent with the *c* = 1 power law associated with the fractal globule, and the former this time suggesting non-universal behaviour. We provide additional details on the fits for other types of simulations in Suppl. Table S2 and Suppl. Fig. S11; we give the same for contacts defined through spatial distance rather than base-pairing in Supplementary Table S3 and Suppl. Fig. S12.

The correlation between simulated and experimental contact maps indicates that our model captures key features of the Zika genome’s 3D organization. Importantly, unlike most RNA secondary-structure algorithms, our approach also allows for pseudoknots and we can therefore analyse their frequency of formation.

The fact that experimental and simulated contact maps (Fig. 1, Fig., Suppl. Fig. S13) contain multiple crossing interactions (specific examples in Suppl. Figs. S14-S15) is already suggestive of pseudoknot formation. However, experimental data alone cannot distinguish whether such crossings originate from a single RNA molecule or represent an average of a population of different RNAs. Our simulations can instead answer this question as we simulate structures for single molecules. Many contacts and therefore many pseudoknots in our simulations are transient. For that reason, we focus on the most common pseudoknots in the simulations with stacking with and without confinement. We do this by first using bpRNA [42] on each simulation frame to parse pseudoknots as the minimum set of base pairs that need to be removed from the arc diagram to yield a pseudoknot-free structure. Next, we identify the most common pseudoknot of four consecutive pairs across all frames separately in unconfined and confined simulations. For the unconfined simulations, this is an alternative conformation to a known set of pseudoknots near the 3′ end of the viral genome. While we cannot independently validate these potential newly identified pseudoknots, we show their secondary structure diagrams in Suppl. Fig. S15 and their frequency alongside diagrams illustrating the experimental support for them. Interestingly, these two most common simulated pseudoknots are supported by about an order of magnitude more reads in COMRADES (in cells) and SPLASH (in virion) experiments than the

We also compared the complexity of pseudoknots across structures in terms of a measure of pseudoknot topology, known as the genus [43, 44] (Table 2). Structures with no pseudoknots have genus 0, the simplest possible pseudoknot - equivalent to the crossing of two arcs in the diagram - has genus 1, and more entangled architectures have higher genera. We find an average genus of ⟨*g*⟩ = 276.4 and ⟨*g*⟩ = 214.7 for the unconfined and confined case respectively, higher than the expected genus of ∼ 140 based on the fit of *g* versus sequence length presented in [41]. We provide another global characterization of the simulated pseudoknots in the form of histograms of their length distribution across all frames in both types of simulations (Suppl. Fig. S16, details in SI Section 7).

**Table 2:** Average genus calculated based on simulations. The (⟨*g*⟩) value for 3D COMRADES simulations with stacking is in line with the value of ∼ 140 expected for an RNA with the length of the Zika genome [41]. Simulations with RNAfold input predict a much lower genus than expected, whereas those with no account for stacking predict a much higher value, largely due to the formation of base-pair ‘mismatches’, i.e., instances where neighbouring bases are paired to disparate regions of the molecule. The column labelled SD_⟨*g*⟩_ gives the standard deviation of the calculated genus. pseudoknots known from the literature ([30], Suppl. Fig. S14).

| Dataset | $\langle g \rangle$ | $SD_{\langle g \rangle}$ |
| --- | --- | --- |
| ViennaRNA RNAPKplex [12] prediction for the KJ776791.2 Zika sequence | 1 | - |
| 3D Vienna, no confinement | 59.4 | 5.7 |
| unconfined 3D COMRADES, no stacking | 1131.9 | 11.8 |
| unconfined 3D COMRADES with stacking | 276.4 | 12.4 |
| confined 3D Vienna | 72.4 | 7.2 |
| confined 3D COMRADES, no stacking | 844.6 | 46.9 |
| confined 3D COMRADES with stacking | 214.7 | 7.8 |

### Folded RNA molecules are glassy and exhibit nematic ordering

Our 3D simulations can also be used to explore the dynamics of folded RNA in different conditions, something which would be extremely challenging to do experimentally. Because base pairing in a long stem typically involves a large energetic gain, we expect the dynamics of contacts to be slow. We quantify these timescales by computing the ensemble-averaged Pearson correlation coefficient between the matrices of base-pairing contacts over the entire simulated time range. As seen in Fig. (e), these are persistent and decay very slowly; they can also be accurately fitted by a stretched exponential. Since a glass is defined as an amorphous system whose relaxation time is longer than the time scale of observation [45], and it is often associated with stretched exponential relaxations [46], these results suggest that the viral RNA exhibits glass-like behaviour. This is further evidenced in the persistent correlations between interbead distances, see Suppl. Fig. S17.

Interestingly, glassiness is more pronounced in unconfined simulations, and spatial confinement inside the virion significantly fluidizes the system, leading to a dramatic decrease of the typical correlation time, by almost an order of magnitude. This suggests that contacts can rearrange much more readily inside the virion than outside it. We interpret this seemingly surprising result by noting that in the absence of spatial confinement the radius of gyration of RNA is much larger, and the bases paired in a stem need to diffuse a significantly larger distance before forming a pair with another base. Conversely, we expect that inside the tight environment of the virion bases can immediately pair again with another partner after the breaking of a pair, corresponding to a lower kinetic barrier for contact rewiring.

The long stems that arise in simulations with stacking also confer local orientational order to the RNA molecule, as strands participating in a stem tend to be parallel to a common axis. We quantify the nematic order via the Maier-Sauper order parameter *S*, which is defined through the average of the angle *θ*_nem_ that defines the orientation of the segments with respect to the nematic director (the average direction of segments) [47]

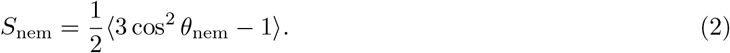

*S*_nem_ is zero if the orientation of all segments is random and unity if they are all aligned in the same direction. We characterize the local nematic order of a bead by taking the average of *S*_nem_ over all beads within a cut-off distnce *l*_cut−off_. Fig. (f) shows the decay of local nematic order ⟨*S*_nem_⟩ with *l*_cut−off_ (see Section 4 in the SI for more details). The decay rate of the approximately exponential decrease of the local order with cut-off defines a characteristic length *l*_nem_ of nematic domains.

In unconfined simulations, *l*_nem_ is 2.99*σ*, whereas in confined ones *l*_nem_ = 2.22*σ*. These results point to a local nematic domain size 2 − 3 nm, which slightly decreases for confined simulations, where stems are more often deformed. That stacking provides the mechanism for local nematic order is confirmed by the smaller values of *l*_nem_ in simulations without stacking (see Suppl. Table S5 and Suppl. Figs. S18-S19).

Nematic domains have been experimentally observed in double-stranded RNA viruses [48], oligonucleotide duplexes [49], and in theoretical studies [50, 51]. We are not aware, however, of any previous reports of nematic ordering in *single-stranded* RNA viruses such as Zika. On the other hand, nematic order likely confers mechanical stability to the viral genome, and it also minimises entanglements, which we speculate will facilitate rearrangement into a functional RNA inside the host cell following capsid disassembly.

### Simulations and experiments demonstrate higher circularization frequency under confinement

It is known that the Zika viral genome circularizes due to strong base-pairing interactions between its 3′ and 5′ ends, and that the circularization state of the viral RNA controls the switching between replication and translation [17, 52]. Circularization in Zika and related viruses such as dengue has been studied extensively via SHAPE-MaP[53–55] - see [17, 56, 57]. It has been suggested that the dengue genome is circular in the virion [56], and evidence from [17] indicates that the Zika genome is linear in vitro, but we do not know of any attempt to quantify the fraction of the circular form. However, using the equivalent NAI-MaP method, Huber et al. [4] measured relatively high reactivities at the bases involved in some circularization contacts, e.g., nucleotides 153-157 in the cyclization sequence [17] (see reactivity scores for the bases involved in circularization in Suppl. Fig. S23). At the same time, chimeric reads from SPLASH from the same work indicate base-pairing in this region (see Figure b). This suggests that cyclization is common in the virion, but coexists with other secondary structures. Therefore, the result from our confined simulations that ≈ 75% of the circularization contacts exist at long times is compatible with existing experimental data on the subject.

To evaluate the effect of confinement on circularization, we first compare the proportions of chimeric reads supporting the 5′ − 3′ interaction in confined and unconfined experimental data. We calculate the sum of proximity ligation *C*_ij_ scores associated with circularization interactions as listed in [17], normalised by the sum of all *C*_ij_ scores in three replicate datasets generated under in vivo conditions and two replicates for experiments with virions. We observe that circularization accounts for 0.14 to 0.52% of the sum of all interaction scores in virions, but only 0.022 − 0.029% in vivo [Fig. (a)]. Thus, confinement increases the relative frequency of circularization by more than an order of magnitude.

Simulations also show that confinement strongly enhances circularisation. Thus, at long times on average ≈ 75% of all circularisation base pairs given in [17] (see the secondary structure diagrams in Fig.) exist in confined simulations, whereas for equivalent unconfined simulations this fraction drops to less than 1% [Fig. (b,c)]. Importantly, using unconfined proximity ligation *C*_ij_ scores as inputs for both confined and unconfined simulations still gives an enhancement of circularisation frequency for confined simulations, showing that confinement genuinely facilitates this interaction (Suppl. Fig. S25). As the enrichment is less than observed in Fig. (d, f), this also suggests that the specific locations and intensities of contacts in experiments with virions favour circularisation.

We characterize the dynamics of circularization by monitoring the fraction of base-pairing contacts shown in the secondary structure diagrams in Fig. (a,b) across all simulation frames after equilibration. Notably, circularization can be temporarily lost in unconfined simulations, whereas it is usually retained throughout the simulation once established in confinement. Note that our simulations also give information of the local 3D structure associated with circularization [Fig. (d)]. As intuitively expected, these involve especially long parallel stems which energetically stabilise the interaction between the two ends; additionally, circularised ends naturally form close to the surface of the sphere, which would facilitate access of proteins to this region once the capsid disassembles in the host cytoplasm.

### Accounting for electrostatic interactions and the capsid protein improves the performance of confined simulations

So far, we have neglected electrostatic interactions. Our justification for this approach is that 1) they are implicit in the experimental proximity ligation scores from which we estimate the base-pairing probabilities and 2) under physiological conditions, both the RNA and the capsid protein within the small volume of the capsid are expected to be ionized, meaning that the local ionic strength is high and electrostatics - highly screened. Nonetheless, we relax this approximation for confined conditions and add a shape-based coarse-grained representation [58] of the capsid protein as a flexible shell based on a crystal structure of the immature Zika virion [59] (see Fig. (h), details in Methods). We find that introducing these additional effects improves the correlation of simulations with experimental data (*R*_Pearson_ = 0.37 vs 0.33 without electrostatics, Table 1). This suggests that in the mature virion, the capsid protein retains at least partial attachment to the outer layers of the virion. This is a valuable insight into the structure of the Zika virus as the capsid protein cannot be imaged in mature virions and previous reports have stated that it may be fully detached from the membrane protein [28].

## Discussion and conclusions

In summary, our analysis of proximity ligation assays reveals striking differences between contact maps obtained in vivo and in virion conditions. First, the probability of contact versus genomic distance (Fig. 1) in virions is consistent with the fractal globule model, which describes a polymer that is confined spatially without being able to equilibrate [32]. Second, the contact probability decay with distance is steeper in vivo, and compatible with an exponent *c* = 3*/*2, which characterises contacts in a random walk, in line with recent predictions that the folding of randomly branched RNA molecule is consistent with that of random walks [34]. Alternative explanations for the different scaling behavior observed in the cytoplasm (in vivo) are possible: for example, active translation in the cytoplasm could periodically unfold RNA, favoring short-range interactions due to faster reannealing kinetics. However, our simulations show that physical constraints are sufficient to explain the differences of scaling behavior.

Because of the length of the Zika genome, our model represents RNA at the level of one bead per nucleotide, and some aspects of the resulting structures are necessarily neglected. For example, base-paired regions in Figures 2-do not adopt A-form helical geometry; the same is true of comparable coarse-grained models of double-stranded DNA in bacteriophages [60], or of dsRNA [50]. The parameters of the model are nevertheless realistic: the stiffness of paired and unpaired RNA and the separation between paired bases are based on literature data (see Methods), yielding models that reproduce experimental contact maps (Table 1) and realistic stem length distributions (Suppl. Fig. S10). We expect our predictions for the global reshaping of RNA base-pairing upon confinement, contact probability relationships, prevalence of pseudoknots and propensity of circularisation to be robust, since these depend on confinement and polymer physics and emerge on scales larger than the radius and pitch of the RNA helix. Nematic order magnitude may instead be affected by helicity, although its presence should be robust as it is due by stacking; it would be of interest to quantify this effect in future work. Approaches that combine coarse-grained simulation with machine learning [61, 62] or atomic resolution Brownian dynamics (ARBD) [63, 64] may also in the future allow atomic detail to be recovered at this scale.

Our coarse-grained molecular dynamics model,(Fig. 2), which uses the experimentally-derived proximity ligation scores as an input, generates contact maps in which the location and intensity of contacts correlate well with the experiment (Fig.). We have also shown that accounting for base-pair stacking in the model is necessary to avoid excessive base pair strain, and the frequent formation of unphysically short stems. Our 3D model predicts a realistic level of pseudoknot complexity, in contrast to models based on ViennaRNA which at most allow for very simple pseudoknots due to computational limitations [65–67]. Such pseudoknots are functionally important [68], for instance as they confer resistance to the action of RNase activity [18] to subgenomic viral RNAs that suppress the host immune response [20].

Our simulations also provide predictions which are outside the reach of current experiments. First, they predict that stacking should promote the formation of small nematic domains, of size 2 − 3 nm, which may help prevent excessive entanglement within the virion. This is in line with recent experimental [48] and theoretical studies [50] that have observed liquid crystalline features in double-stranded RNA. Second, our simulations predict slow and glass-like dynamics of RNA contacts, which may explain why the RNA does not thermodynamically fully equilibrate and hence folds into a fractal globule inside the virion.

Finally, we have observed that circularization contacts, which play an important role in the lifecycle of the Zika virus, occur at a much higher frequency within virions than under in vivo conditions (Fig.). This result is supported by both experiments and simulations, and is functionally important as these contacts determine the balance between transcription and translation once viral particles are released into the host. Importantly, our simulations show that the enhancement of circularization is due in part purely to spatial confinement, and in part due to specific pairing of stretches of nucleotides at the molecule ends, which promote the formation of long stems. These contacts localise close to the surface of the confining capsid, which may facilitate access from RNA-binding proteins once the RNA is in the cytoplasm.

Our approach complements chemical probing methods such as SHAPE [54, 55, 69] that differentiate single- and double-stranded RNA regions by their reactivity, and in the case of mutational profiling with dimethyl sulphate coupled with DREEM clustering [70], quantify the abundance of different secondary structures. Computational modelling techniques such as those we develop here provide additional insight into the dynamics of RNA secondary structure remodelling.

The wealth of data accumulated through proximity ligation studies has not to date been integrated with computational modelling, largely because of the computational cost of applying fine-grained models to long RNAs [71]. Another challenge of interpreting proximity ligation data sets such as those we leverage here is that they are known to contain bias due to the affinity of cross-linking agents to particular double-stranded regions [71, 72].

We have developed a simulation technique which sheds light into both the frequency of contact and the exact contact patterns in our simulations of the Zika viral genome under both in vivo and in virion conditions. We highlight that our approach to studying RNA folding in 3D with and without confinement will be broadly applicable to the study of the spatial organization of RNA in different scenarios. For instance, we could use them to study mRNA vaccines, which are typically delivered in confined carriers such as lipid nanoparticles. As demonstrated in [73], encapsulation within particles in this size range can not only promote intramolecular RNA contacts, but also stabilize tertiary folding and thereby promote RNA activity. Our methods can also be adapted to study nucleic acids under confinement in a broader range of contexts, for example, non-coding RNAs in cellular structures such as stress granules, DNA in adeno-associated viruses used for gene therapy delivery, and other problems.

## 1 Methods

### 3D Vienna

We use the RNAfold program from the ViennaRNA package, version 2.6.4 [12], as input for one type of simulation. Instead of taking the mininum-free-energy secondary structure for the viral RNAs, we use the base-pairing probabilities for the thermodynamic ensemble for the probabilities *p*_ij_ of forming a pair between bases ‘i’ and ‘j’ that are within *d*_thr_ of each other. Choosing an ensemble-based approach rather than one focussed on individual structures is appropriate as the frequency of even the minimum-free-energy structure can be vanishingly small [74] - this is estimated via RNAfold to be of the order of 10^−0.01*L*^ for randomly permuted sequences [40], which for the Zika genome yields ∼10^−108^. Note that even though ViennaRNA explicitly disallows structures with pseudoknots due to dynamic programming constraints [12], the base-pairing patterns that arise in our simulations can potentially contain pseudoknots as the ensemble base-pairing probabilities for bases within different stem loops are non-zero. As RNAfold already accounts for stacking, we do not use our own approximate model for that (see below) and set Δ*G*_penalty_ = Δ*G*_stacking NNN_ = Δ*G*_stacking NN_ = 0.

### 3D COMRADES

As an alternative to using RNAfold predictions for the base-pairing probabilities *p*_ij_, we incorporate experimental data from proximity ligation assays into the model. Contact maps predicted with RNAfold have a much lower density of contacts than that measured experimentally under both in vivo [1] and in virion conditions [4]. These differences may be due to a variety of factors, including RNA-protein interactions that may reduce electrostatic self-repulsion, especially within the nucleocapsid, as well as spatial confinement in the virion. We take the experimentally determined proximity ligation (COMRADES) scores, which heuristically measure the strength of pairing between every two bases [1]. For the pair between bases ‘i’ and ‘j’, the proximity ligation score *C*_ij_ is defined as the number of chimeric reads that contain a pair between these two bases for which UNAFold predicts an i-j base pair in the minimum free energy structure [1]. We use sets of proximity ligation scores calculated based on aggregate data from three replicate COMRADES experiments for in vivo conditions [1] and from two SPLASH replicates with virions [4].

As the large majority of base pairs are rarely observed, *C*_ij_ ∼ 1, instead of using all proximity ligation scores, we focus on the ones with the highest *C*_ij_, taking the top 5% of them. We calculate the cumulative distribution function (CDF) for these top ones, and set the base-pairing scores for nucleotides ‘i’ and ‘j’ (*p*_ij_) to the value of the CDF for that score; for scores outside the top 5% bracket under the given conditions, we use *p*_ij_ = 0; we also set *p*_ij_ = 0 for beads that are beyond a distance *d*_thr_ of each other. In contrast to the ones obtained from RNAfold, the base-pairing scores calculated in this way are not probabilities as the inequality ∑_*j*_ *p*_ij_ ≤ 1 does not necessarily hold here since one base may form pairs with top-ranking *C*_ij_ base-pairing scores with multiple others.

In both 3D Vienna and 3D COMRADES, we treat *p*_ij_ as the probability that two bases will pair if they are within *d*_thr_ of each other. Since we perform pairwise checks that go through both ‘i’ and ‘j’, there are two opportunities for each pair to form, which means that the probability of forming a pair in these two trials obeys the equation *p*_ij_ = *w*_ij_ + *w*_ij_(1 − *w*_ij_). For that reason, we set the probability of forming a base pair at every step to 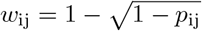 and the probability of breakage to 1 - *w*_ij_.

### Models for stacking

We account for the energy gain of stacking base pairs by modifying the probability of forming a base pair between beads that are within *d*_thr on_ of each other.

Specifically, the stacking model stipulates that, if one or both of the nearest neighbours of the potential base pair under consideration are paired so that forming the potential base pair will extend a stem, this results in an energy gain that increases the probability of forming this base pair. We calculate this gain using an estimate of the free energy change due to stacking and pairing of distant bases that contains additive contributions corresponding to the state of the neighbouring bases. We then multiply the base-pairing score *w*_ij_ by a Boltzmann factor, *w*_ij stacked_ = min(*w*_ij_ exp{−Δ*G*_stacking_*/*(*k*_B_*T*)}, 1). The contributions are:

a. *Stacking with nearest neighbours*. The energy gain is Δ*G*_stacking NN_ = − *k*_B_*T* in the presence of one nearest neighbour, and twice that if both neigbours in the stem are present.
b. *Stacking with next-nearest neighbours*. If both a nearest neighbour and the associated next-nearest neighbours exist, we add another stacking contribution Δ*G*_stacking NNN_ = Δ*G*_stacking NN_*/*2 = − *k*_B_*T/*2.
c. *Neighbouring bases paired to non-adjacent bases*. In this case, we add a positive contribution to the energy, Δ*G*_penalty_ = (3*/*2)*k*_B_*T*, as this should be disfavoured by stacking.

In all cases, when stochastically breaking base pairs, we set the probability of breaking to *w*_ij break stacked_ = 1 − *w*_ij stacked_.

### Simulations with electrostatic interactions

Electrostatic interactions are tractable for the case of RNA within virions. The Zika viral genome experiences non-specific electrostatic attraction to the positively charged capsid (C) protein [28]. During the maturation of the Zika virus, the transmembrane (TM) region of the precursor membrane (prM) protein rearranges, causing C dimers to bind to one prM-TM regions in the mature virion as opposed to three in the immature form [28]. Cryo-EM studies cannot resolve the C protein in the mature virion as it may be completely or partially detached from the TM region of the mature membrane (M) protein, or it may remain bound to one M-TM, which would give it three possible positions within every asymmetric unit, making it more difficult to fit its position within the virion [28]. Here, we model the RNA-C interactions on the basis of a high-resolution cryo-EM structure of the capsid of the immature virion [28], creating a shape-based coarse-grained (SBCG) representation of it. This corresponds to the approximation that the inner capsid of the mature virion has a flexible but well-defined structure that is attached to the M protein and confines the viral genome. The SBCG method [58] mimics the shape of a crystal structure by representing it as a set number of beads connected by pairwise and angle bonds between beads separated by less than 8 nm = 10*σ*. We apply this approach to the structure [59] from [28], using one bead per C monomer. Both types of bonds between SBCG beads are harmonic, with the energy of the pairwise bonds following Eq. (S6) and that of the angle bonds being *E*_harmonic angle_ = *K*_*θ*_ (*θ* − *θ*_0_)^2^. The individual equilibrium length *r*_0_ and angle *θ*_0_ for each bond is optimized by the SBCG algorithm for maintaining the shape of the crystal structure, and the values for the spring constants are set to *K* = 20 and *K*_*θ*_ = 20 for all interactions. We account for the excluded-volume interactions between beads ‘i’ and ‘j’ belonging to different types via the Weeks-Chandler-Anderson potential, Eq. (S5) in which the interaction energy parameter *ϵ* and bead diameter *σ* are substituted with parameters that depend on the bead types in the i-j pair, *ϵ*_ij_, *σ*_ij_.

These are determined from arithmetic mixing rules, 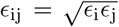 and 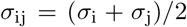, where *ϵ*_i_ is the interaction energy between beads of type ‘i’, and *σ*_i_ is the bead diameter of that type. For nucleotide beads, we use *σ*_RNA_ = 1 as before; for sodium ions, we set *σ*_Na_+ = 0.6*σ*, approximating the hydrated ionic diameter; for all SBCG bead types we set *σ*_SBCG_ to the approximate diameter determined by the SBCG algorithm as the gyration radius of the group of atoms represented by the bead. We set the dimensionless mass of nucleotide beads to 1 and use the approximate molar mass of a single nucleotide, 320 g/mol, to calculate the dimensionless masses of all other beads. We use the same interaction energy as before for RNA and Na^+^, *ϵ* = *k*_B_*T*, and set *ϵ*_SBCG_ = 4*k*_B_*T* for all other beads. Bead masses and positions are assigned in correspondence with the atomic masses represented as the given bead type. We use Isoelectric point calculator [75] to calculate that under cytoplasmic conditions (pH = 7.4) every bead carries a charge of ≈ 20*e*, where *e* is the elementary charge. In addition to the other interactions described so far, we include a Coulomb potential,

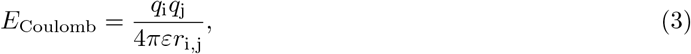

where the charges *q*_i_, *q*_j_ are expressed in SI units and *ε* = 75*ε*_0_ is the approximate dielectric permittivity of water at 310.15 K and *ε*_0_ is that of vacuum. We position the confining potential at *r* = 18.75*σ* to reflect the outer radius of the capsid layer in the cryo-EM serving as the basis for our description [59]. We take the final frames of 20 confined simulations with stacking as the initial conditions for simulations involving electrostatics, an SBCG representation of the capsid protein. In each simulation, we add a number of univalent cations that, together with the C protein, exactly neutralizes the charge of the RNA - in the absence of data on the electrolyte content of the virion, we assume that the capsid together with the RNA and hydrated sodium ions form an electrically neutral system. We generate random initial positions of the cations that do not overlap with RNA beads and are within the confining sphere with radius 16.4*σ* containing the RNA beads from the simulation without electrostatics; we use the same SBCG representation of the capsid in all simulations. As with simulations without electrostatics, we run several equilibration stages. In the first, the RNA is equilibrated the RNA with stiff harmonic and angle bonds combined with soft repulsion between beads. This is followed by switching to the bond and angle styles for the main simulation, removing the base-pairs from the initial configuration, switching to excluded volume interactions based on Eq. (S5), switching electrostatic interactions on and ramping up the absolute value of the charges from 0 to their correct values for all beads. During these equilibration stages we limit the maximum displacement of atoms to avoid numerical issues. Once all interactions are set to their correct values for the main simulation, we run the simulation for an additional 50*τ* before allowing base-pairs to form. The main simulations run for 6 × 10^3^*τ* units, of which we use the last 5 × 10^3^*τ* in our analysis; the long-range electrostatic interactions require a shorter time step then before, Δ*t* = 10^−3^*τ*. As in the simulations with no electrostatics, we update the list of base pairs every 10*τ*.

### Topological characterization of RNA structures

Pseudoknots can be classified based on the value of a topological characteristic called the *genus* [43, 44]. For an arc diagram of RNA secondary structure, the genus is defined as follows [44]. First, one joins the 3′ and the 5′ ends into a circle. Then, one draws the lines representing base pairs on the outside of the circle. The genus is then the minimum number of handles that one needs to carve in a punctured sphere in order to draw the graph on it without any crossings.

It is convenient to represent the arc diagram as a permutation (*π*) in which each position from one to the length of the RNA contains the index of the base that this position is paired to [44]. A closed loop (cycle) in this permutation is a sequence of elements in which the last element maps back to the first [76]. The genus *g* of an RNA arc diagram is simply related to the number of base-pairings *p* in the arc diagram and the number of closed loops *l* that characterize the composite permutation *ϕ · π*, where *ϕ* = (2, 3, 4, …, *L*, 1) is the cyclic right-shift permutation [44]:

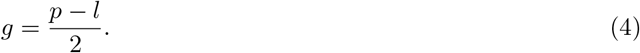

Arcs that are parallel to each other and intersect the same other arcs are equivalent to each other. Any sheaf of these arcs can be collapsed into a single renormalized arc without affecting the genus because removing an equivalent arc decreases *l* and *p* by one, keeping their difference and hence *g* constant, (4). We provide arc diagrams simplified in this way in Suppl. Fig. S13.

## Supplementary information

We provide additional details on processing experimental and simulated data in the Supplementary Information along with additional figures, code, movies illustrating several types of simulations and data on base-pairing patterns in all post-equilibration frames for our simulations with stacking (https://figshare.com/s/9c24ece7518d20129ac5).

## Author contributions

J.K.N.: conceptualization, methodology, software, data curation, formal analysis, investigation, writing—original draft and writing—review and editing; J.Y.L.: data curation, formal analysis D.M.: conceptualization, methodology, formal analysis, software, writing—review and editing G.K.: conceptualization, methodology, writing—review and editing.

## Acknowledgements

J.K.N. acknowledges funding from the MRC (Grant MC FE 00035) Cross Disciplinary Fellowship (XDF) Programme. We thank Dr Elly Gaunt (Roslin Institute, University of Edinburgh) for helpful discussions.

## Footnotes

1 Note though that for contact between internal points the value of *c* changes to *c ≈* 2 [33].

## Notes

### Competing Interest Statement

The authors have declared no competing interest.

### Summary of Updates

The text and some of the methods have been updated; the new version includes the role of electrostatic interactions.

