## Supplementary Material for "3D confinement reshapes RNA folding and enhances circularisation in the Zika virus"

### Supplementary Information to the article 3D confinement reshapes RNA folding and enhances circularisation in the Zika virus

J. K. Novev*[a], J. Y. Lau[a], D. Marenduzzo[b], and G. Kudla[a]

[a]Institute of Genetics and Cancer, University of Edinburgh, Western General Hospital, Crewe Road South, Edinburgh EH4 2XU, UK

[b]School of Physics and Astronomy, University of Edinburgh, James Clerk Maxwell Building, Peter Guthrie Tait Road, Edinburgh EH9 3FD, UK

#### 1 Processing experimental data

We disregard the difference between counts and $C_{\mathrm{ij}}$ base-pairing scores (COMRADES scores) for chimeras ligated in $5' - 3'$ and $3' - 5'$ orientations, summing the contributions for both types.

##### Coarse-graining of contact maps

We coarse-grain the matrices that contain information on the number of experimentally observed counts supporting a contact or the calculated probabilities of pairing between beads from our simulations. In each of these matrices, element $p_{\mathrm{ij}}^{\mathrm{sim}}$ refers to base 'i' pairing with base 'j'. We coarse-grain the data from these matrices by binning it into non-overlapping bins of size $n_{\mathrm{CG}}$. As $L$ is not typically divisible by $n_{\mathrm{CG}}$, the final bins are smaller than the rest. Except otherwise noted, we coarse-grain with $n_{\mathrm{CG}} = 30$ as this is the typical length of each arm of the chimeras sequenced in the proximity ligation experiments.

###### Binning of experimental maps

We sum the counts recorded for every coarse-graining window with $\mathrm{i} \in [\mathrm{i}_{\mathrm{min}}, \mathrm{i}_{\mathrm{max}}]$ and $\mathrm{j} \in [\mathrm{j}_{\mathrm{min}}, \mathrm{j}_{\mathrm{max}}]$.

###### Coarse-graining simulated distance maps

te the centre of mass of all beads in each window; within a window, the bead index 'i' runs from $\mathrm{i}_{\mathrm{min}}$ to $\mathrm{i}_{\mathrm{max}}$. For each processed frame, we calculate the matrix of distances $d_{\mathrm{IJ}}^{\mathrm{sim}}$ between the centres-of-mass of coarse-grained sets of beads with coarse-grained indices 'I' and 'J' equal to the integer part of the ratio between the lower bound of the respective interval and the coarse-graining size $n_{\mathrm{CG}}$, $\mathrm{I} = \mathrm{floor}(\mathrm{i}_{\mathrm{min}}/n_{\mathrm{CG}})$ and $\mathrm{J} = \mathrm{floor}(\mathrm{j}_{\mathrm{min}}/n_{\mathrm{CG}})$; 'I' and 'J' both lie in the range $[0, \mathrm{ceil}(L/n_{\mathrm{CG}})]$. We then set the contact matrix $C_{\mathrm{IJ}\,\mathrm{dist}}^{\mathrm{sim}}$ to 1 for all sets I, J for which $d_{\mathrm{IJ}}^{\mathrm{sim}} < d_{\mathrm{cont}} = 10\sigma$. We then average the matrix $C_{\mathrm{IJ}\,\mathrm{dist}}^{\mathrm{sim}}$ over all processed frames and runs in an ensemble.

###### Coarse-graining simulated base-pairing probability maps

We calculate the sum of base-pairing probabilities $p_{\mathrm{window}}^{\mathrm{sim}}$ for every coarse-graining window in which $\mathrm{i} \in [\mathrm{i}_{\mathrm{min}}, \mathrm{i}_{\mathrm{max}}]$ and $\mathrm{j} \in [\mathrm{j}_{\mathrm{min}}, \mathrm{j}_{\mathrm{max}}]$. We then normalize this sum of probabilities by the maximum value $p_{\mathrm{window,max}}^{\mathrm{sim}}$ that it can take if all bases are fully paired. This differs depending on the type of window in question.

1. For windows in which there are no elements in common between the sets $[\mathrm{i_{min}}, \mathrm{i_{max}}]$ and $\mathrm{j} \in [\mathrm{j_{min}}, \mathrm{j_{max}}]$, $p^{\mathrm{sim}}_{\mathrm{window,max}}$ is equal to the number of elements in the smaller set, $p^{\mathrm{sim}}_{\mathrm{window,max}} = \min(\mathrm{i_{max}}-\mathrm{i_{min}}, \mathrm{j_{max}}-\mathrm{j_{min}})+1$.

2. For windows in which the bases covered by both indices are identical, $[\mathrm{i_{min}}, \mathrm{i_{max}}] \equiv \mathrm{j} \in [\mathrm{j_{min}}, \mathrm{j_{max}}]$, the maximum pairing probability is equal to the number of bases in the window divided by two, $p^{\mathrm{sim}}_{\mathrm{window,max}} = (\mathrm{i_{max}} - \mathrm{i_{min}} + 1)/2$. This comes from the condition that the pairing probabilities for each base 'i' add up to no more than one, $\sum_{\mathrm{j}\in[\mathrm{j_{min}},\mathrm{j_{max}}],\ \mathrm{i<j}} p_{\mathrm{ij}} \leq 1$. Summing these identities for $\mathrm{i} \in [\mathrm{i_{min}}, \mathrm{i_{max}}]$, we get $\sum_{\mathrm{i}\in[\mathrm{i_{min}},\mathrm{i_{max}}]} \sum_{\mathrm{j}\in[\mathrm{j_{min}},\mathrm{j_{max}}],\ \mathrm{i<j}} p_{\mathrm{ij}} \leq \mathrm{i_{max}} - \mathrm{i_{min}} + 1$. Each of the base-pairing probabilities appears twice in the double sum because it is subject to the condition that pairing probabilities add up to no more than 1 for both participating bases, meaning that $2\sum_{\mathrm{i}<j} p_{\mathrm{ij}} \leq \mathrm{i_{max}} - \mathrm{i_{min}} + 1$. As the matrix of base-pairing probabilities is symmetric ($p^{\mathrm{sim}}_{\mathrm{ij}} = p^{\mathrm{sim}}_{\mathrm{ji}}$) and also contains elements with $\mathrm{j} > \mathrm{i}$, the correct normalization is $p^{\mathrm{sim}}_{\mathrm{window,max}} = \mathrm{i_{max}} - \mathrm{i_{min}} + 1$. In practice, we minimize memory use by performing most manipulations with the lower triangular matrix $P^{\mathrm{sim}}_{\mathrm{ij}}$, which we derive from $p^{\mathrm{sim}}_{\mathrm{ij}}$ by setting elements with $\mathrm{j} > \mathrm{i}$ to zero; in this case, the normalization is $P^{\mathrm{sim}}_{\mathrm{window,max}} = (\mathrm{i_{max}} - \mathrm{i_{min}} + 1)/2$.

We then calculate the average base-pairing probability for the window and assign it to the respective element of the coarse-grained matrix $p^{\mathrm{CG}}_{\mathrm{IJ}} = p^{\mathrm{sim}}_{\mathrm{window}}/p^{\mathrm{sim}}_{\mathrm{window,max}}$, with coarse-grained indices 'I' and 'J' equivalent to those for the distance-based contact matrix.

#### Symmetrizing experimental count matrices and calculating simulated base-pairing probabilities from simulations and secondary structure prediction algorithms

We symmetrize matrices of experimental counts $N_{\mathrm{ij}}$. We do this by adding all elements $N_{\mathrm{ij}}$ above the matrix diagonal, for which $\mathrm{i} < \mathrm{j}$, to the corresponding element below the diagonal, $N_{\mathrm{ji}}$, and set $N_{\mathrm{ji}} = 0$. Finally, we copy all elements below the diagonal above the diagonal so that $N_{\mathrm{ji}} = N_{\mathrm{ij}}$.

We calculate simulated contact probability matrices by counting the number of frames that contain a pair between bases 'i' and 'j' in an ensemble of simulations $N_{\mathrm{frames\ pair\ ij}}$, then averaging it over time and ensemble dividing by the number of frames analyzed and the number of runs in the ensemble, $p^{\mathrm{sim}}_{\mathrm{ij}} = N_{\mathrm{frames\ pair\ ij}}/(N_{\mathrm{frames}}N_{\mathrm{runs}})$. These matrices are symmetric by definition.

#### Contact probability exponents

For experimental data, we calculate the contact probability as a function of the separation between segments $p_{\mathrm{contact\ exp}}(l)$ as follows. We first find $N_{\mathrm{contacts\ exp}}(l)$, the sum of the number of chimeras supporting contacts between non-overlapping bins of bases with width $l_{\mathrm{bin}}$, then normalize this sum by the overall number of contacts,

$$p_{\mathrm{contact\ exp}}(l) = N_{\mathrm{contacts\ exp}}(l)/\sum_{l} N_{\mathrm{contacts\ exp}}(l). \tag{S1}$$

For simulated data, we calculate two types of contact probabilities for our simulations based on two ways to define a contact.

1) We calculate the distances $d_{\mathrm{interbin\ ij}}$ between the centres-of-mass of the beads binned into bins i and j, both of width $l_{\mathrm{bin}}$. For each simulation frame, we record a contact wherever $d_{\mathrm{interbin\ ij}} < d_{\mathrm{contact}} = 10\sigma$. Doing this for all frames yields the number of distance-based contacts recorded for each bin, $N_{\mathrm{contacts\ distance\ ij}}$, from which we calculate a contact probability $p_{\mathrm{contact\ distance\ ij}}$ for each ij window via dividing by the number of processed frames and the number of simulations in the ensemble. Then, for each separation $l$, we calculate the contact probability as the sum of values of $p_{\mathrm{contact\ (distance)\ ij}}$ for that separation for which $l_{\mathrm{bin}}|\mathrm{i}-\mathrm{j}| = l$, normalized by the sum of all $p_{\mathrm{contact\ (distance)\ ij}}$,

$$p_{\mathrm{contact\ distance}}(l) = \frac{\sum_{l_{\mathrm{bin}}|\mathrm{i-j}|=l} p_{\mathrm{contact\ (distance)\ ij}}}{\sum_{\mathrm{i,j}} p_{\mathrm{contact\ (distance)\ ij}}}. \tag{S2}$$

2) We calculate a contact probability defined through base-pairing in the simulation as

$$p_{\text{contact bp}}(l) = \frac{\sum_{l_{\text{bin}}|\text{i}-\text{j}|=l} p_{\text{contact bp ij}}}{\sum_{\text{i,j}} p_{\text{contact bp ij}}} \quad \text{(S3)}$$

where the only difference with (S2) is that we use the number of base pairs for the window indexed ij, rather than the number of distance-based contacts for the same window.

##### Data from secondary structure prediction software

We use predictions for the base-pairing probabilities in the thermodynamic ensemble from RNAfold [1], UNAfold [2] and LinearPartition [3], then treat those in the same way as $p_{\text{contact bp ij}}$ in (S3).

#### Correlations between contact maps

We flatten the symmetrized, coarse-grained matrices with experimental counts, predictions from secondary structure algorithms and ensemble- and time-averaged simulated base-pairing probabilities. We then characterize the pairwise correlations by calculating the respective Pearson ($R_{\text{Pearson}}$) and Spearman ($R_{\text{Spearman}}$) correlation coefficients. All contact-map correlations are highly significant statistically.

#### Exponent fitting

Eq. (1) can only be expected to apply to intermediate genomic distances $l$ for which the length of the corresponding section of the RNA fibre is greater than the RNA persistence length $l_{\text{p}}$, but smaller than the genome length $L$.Moreover, experimental data from proximity ligation assays does not accurately reflect contact probabilities for $l \sim 1$ because such contacts correspond to chimeric reads that map to a contiguous region of the viral genome. Such reads are discarded in the processing of COMRADES and SPLASH because it is not possible to distinguish whether they come from a pair of segments that were fragmented and subsequently ligated or a single long fragment, see Gabryelska et al. for more on the subject [4]. In simulations, the permanent bonds between consecutive beads mean that the latter are always in contact and do not follow the limiting law in (1), either.

We fit the various types of $p_{\text{contact}}(l)$ data to the power law (1) for intermediate values of $l$, typically taking only values $l < l_{\text{cut-off}}$, where $l_{\text{cut-off}}$ is the minimum value at which $p_{\text{contact}} = 0$. We provide slopes, $R^2$ values, and fitting ranges for experiments and secondary structure prediction algorithms in Supplementary Table S1, for contacts defined through distance in Supplementary Table S3 and for contacts defined through base-pairing in Supplementary Table S2.

#### Sequences

We study the Asian strain of the Zika virus, using sequence KX197192.1 (isolate ZIKV/H.sapiens/Brazil/PE243/2015) [5] for our simulations under unconfined conditions (simulating COMRADES in vivo measurements from [6]), and sequence KJ776791.2 (French Polynesia)[7] for confined conditions (simulating SPLASH measurements in virions from [8]). These two sequences differ by $\sim 20$ single-nucleotide substitutions and are 99.76 % identical.

#### Additional experimental data

We plot the contact map, contact probability vs. distance and distribution of interaction lengths for a number of additional proximity ligation datasets for various strains of the Zika virus and the Dengue 1 virus. We process SPLASH datasets from experiments performed in the Wan lab (Genome Institute of Singapore, A*STAR) [8] with both cells and virions for 1) the African strain of the Zika virus (sequence AY632535.2), 2) the Brazil strain of the Zika virus (sequence KU497555.1), 3) the Singapore strain of the Zika virus (sequence KY241702.1), and 4) the Dengue 1 virus (sequence EU081230.1). We process a dataset from the same source

[8] that complements the data for virions we discuss in the main text: this data is for the French Polynesian strain (sequence KJ776791.2) and is obtained from cells. Additionally, we analyze data from experiments with cells from [9] for 1) the Puerto Rico strain (sequence KX377337.1) and 2) the Uganda strain (sequence KX377335.1); for the Puerto Rico strain, we also analyze a dataset for in vitro conditions. Of these, the Puerto Rico strain is most closely related to the Brazil (KX197192.1) and French Polynesia (KJ776791.2) strains we study in the main text, differing only by point substitutions and having 99.76% sequence identity to KJ776791.2. We compare specific interactions in these three strains in Suppl. Fig. S1. We process all data via the hyb2 bioinformatics pipeline developed by Lau et al. [10].

### 2 Coarse-grained molecular dynamics

We construct a coarse-grained molecular dynamics model of the viral genome within the software package LAMMPS (Large-scale Atomic/Molecular Massively Parallel Simulator, 17th April 2024) [11]. We use one coarse-grained bead per nucleotide for the viral RNA, with all four types of nucleotides represented in the same way, and simulate the evolution of both unconfined and confined viral genomes via Brownian dynamics.

For the $3'-5'$ phosphodiester bonds between consecutive nucleotides, we use a finite extensible nonlinear elastic (FENE) potential [12] according to which the interaction energy between two bonded beads 'i' and 'j' that are separated by a distance of $r_{\mathrm{i,j}} = |\boldsymbol{x}_{\mathrm{i}} - \boldsymbol{x}_{\mathrm{j}}|$, where $\boldsymbol{x}_{\mathrm{i}}$ is the position vector of bead 'i', is

$$E_{\mathrm{FENE}} = -\frac{1}{2} K_{\mathrm{FENE}} R_0^2 \ln \left[ 1 - \left( \frac{r_{\mathrm{i,j}}}{R_0} \right)^2 \right], \tag{S4}$$

where $\sigma$ is the bead diameter, $K_{\mathrm{FENE}} = 30\, k_{\mathrm{B}}T/\sigma^2$ is a constant characterizing the bond strength, and $R_0 = 1.6\sigma$ is the maximum extent of the bond.

In addition, we model the excluded-volume interaction between any two beads 'i' and 'j' with the purely repulsive Weeks-Chandler-Anderson potential:

$$E_{\mathrm{WCA}} = \begin{cases} 4\epsilon \left[ \left( \frac{\sigma}{r_{\mathrm{i,j}}} \right)^{12} - \left( \frac{\sigma}{r_{\mathrm{i,j}}} \right)^{6} + \frac{1}{4} \right] & r_{\mathrm{ij}} < 2^{\frac{1}{6}} \sigma, \\ 0 & \text{otherwise.} \end{cases} \tag{S5}$$

In our simulations, we set $\epsilon = k_{\mathrm{B}}T$, where $k_{\mathrm{B}}$ is the Boltzmann constant and $T$ is the temperature.

We model the hydrogen bonds in the viral genome with a harmonic potential,

$$E_{\mathrm{harmonic}} = K \left( r - r_0 \right)^2, \tag{S6}$$

with $K = 1$ and $r_0 = 2.4\sigma$ being the spring constant and equilibrium bond length, respectively. We allow for base pairs to form and break dynamically, with the probability of these two events given by RNAfold base-pairing probabilities or by experimental data on base-pairing, see below for details. We update the list of paired bases every $2 \times 10^3$ time steps, allowing base pairing for bases at a distance $r < d_{\mathrm{thr}} = 10\sigma$ of each other. For each unpaired base 'i', we calculate what bases are within $d_{\mathrm{thr}}$, pick a base 'j' at random from these, then attempt to stochastically form a pair between them with probability $p_{\mathrm{ij}}$. If a base pair is not formed, we continue the process until we exhaust all possible bases 'j' that are within $d_{\mathrm{thr}}$, or form a pair. For each already existing base pair between bases 'j' and 'k', we stochastically attempt to break it with probability $1 - p_{\mathrm{jk}}$.

We account for the bending stiffness of the viral RNA by including a potential that depends on the angle $\theta$ between the bond vectors of two consecutive bonds, $\cos\theta = \boldsymbol{r}_{\mathrm{i+1,i}} \cdot \boldsymbol{r}_{\mathrm{i,i-1}} \left( |\boldsymbol{r}_{\mathrm{i+1,i}}||\boldsymbol{r}_{\mathrm{i,i-1}}| \right)^{-1}$, with the bond vectors defined as $\boldsymbol{r}_{\mathrm{i,j}} = \boldsymbol{x}_{\mathrm{i}} - \boldsymbol{x}_{\mathrm{j}}$. The angle bond potential we use is

$$E_{\mathrm{angle}} = \frac{k_{\mathrm{B}} T l_{\mathrm{p}}}{\sigma} \left( 1 + \cos\theta \right), \tag{S7}$$

where $l_{\mathrm{p}}$ is the persistence length of the RNA fibre. Literature data indicates that single-stranded RNA is much more flexible than double-stranded RNA, and to capture that, we set the persistence length $l_{\mathrm{p}}$ to

$\sim 1\,\mathrm{nm}$ [13] for angle bonds for which one or fewer of the beads defining the angle is part of a base pair, and $\sim 60\,\mathrm{nm}$ [14] otherwise.

We run ensembles of simulations with identical input parameters and different random initializations of the polymer chain as a random walk. We run 100 simulations with no electrostatics, $C_{\mathrm{ij}}$ proximity ligation score input with and without stacking for both the unconfined and the confined case. For all other types of simulations, we use an ensemble size of 20. Every simulation contains two equilibration stages that remove possible overlaps between the beads due to the random initialization of their coordinates and allow for relaxation of the RNA molecule in the absence of base pairing. In the first stage, lasting $10^3\tau$, the covalent bonds during the beads use the harmonic potential (S6) with $K = 100 k_{\mathrm{B}}T/\sigma^2$ and $r_0 = 1.1\sigma$, and the persistence length is $l_{\mathrm{p}} = 10\sigma$ for the angle bonds. In the second stage, also with duration $10^3\tau$, the covalent bonding potentials between beads are switched to FENE, (S4), and the persistence length is set to 1 nm. Unless otherwise noted, the duration of the simulations of RNA base-pairing is $10^4\tau$, with a time step of $5 \times 10^{-3}\tau$; we save the output from the simulation every $2 \times 10^4$ time steps, which yields 100 frames overall.

We choose a mapping between the model and experimental data that approximately preserves the volume occupied by each nucleotide. This volume can be estimated by the volume of a cylinder of radius $R \sim 0.5\,\mathrm{nm}$ and height $h \sim 0.33\,\mathrm{nm}$, $V_{\mathrm{nucleotide}} \approx \pi R^2 h$, with dimensions based on typical values for B-DNA, see e.g. Ref. [15] and the associated PDB entry (1ZEW). Equating this to the volume of a spherical bead with diameter $\sigma$, we obtain $\sigma \approx 0.8\,\mathrm{nm}$. For simulations with confinement, we introduce a harmonic potential that restricts the viral genome to a spherical region of radius $R_{\mathrm{conf}}(t)$, with $t$ denoting time:

$$R_{\mathrm{conf}}(t) = \begin{cases} R^0_{\mathrm{conf}} + 2\frac{t}{t_{\mathrm{sim}}}\left(R^{\mathrm{fin}}_{\mathrm{conf}} - R^0_{\mathrm{conf}}\right) & t < t_{\mathrm{sim}}/2, \\ R^{\mathrm{fin}}_{\mathrm{conf}} & \text{otherwise.} \end{cases} \tag{S8}$$

We set the initial confinement radius $R^0_{\mathrm{conf}}$ to a value much greater than the typical gyration radius of the RNA molecule, and for the final confinement radius $R^{\mathrm{fin}}_{\mathrm{conf}}$ we choose $R_{\mathrm{capsid\ inner}} = 13\,\mathrm{nm}$, which is the approximate radius of the inner capsid of the Zika virus [16]. Given the length of the viral genome, $L = 10807$ nt [6], this corresponds to an RNA volume fraction of $\approx 31\%$ in the inner capsid, and a viral genome density of $\approx 1170$ Mb/µm$^3$, more than 50-fold higher than the typical genome density in the nucleus of a diploid human cell [17].

### 3 Simulation time scale

Base pairing and other RNA properties such as the molecular gyration radius equilibrate on a time scale of $\sim 50$ frames for unconfined simulations ($5 \times 10^3\tau$). For simulations with confinement, the time-dependence of the confining potential, (S8), influences the time scale for equilibration of base-pairing, which is why we ran additional simulations with a larger number of frames (200 vs 100 time frames for the unconfined case). The simulation time unit $\tau$ maps to the diffusional time scale $\sigma^2/D$, where $D$ is the diffusion coefficient for a bead. From the Einstein-Smoluchowski equation, the diffusion coefficient is $D \sim k_{\mathrm{B}}T/(\eta\sigma)$, giving $\tau = \sigma^3\eta/(k_{\mathrm{B}}T)$, with $\eta$ being the viscosity of the medium, $\eta \sim 10^{-3}$ Pa s. This gives $\tau \approx 0.12$ ns and $10^4\tau = 1.2$ µs for the typical duration of our simulations. We save the simulation output at intervals of $2 \times 10^4$ time steps, which means that frames are separated by $10^2\tau$.

### 4 Nematic order calculation

For each bead 'i' in the chain, we calculate a tangent vector:

$$\boldsymbol{t}_{\mathrm{i}} = \begin{cases} (\boldsymbol{r}_{\mathrm{i}+1} - \boldsymbol{r}_{\mathrm{i}}) & \text{if } \mathrm{i} = 1; \\ (\boldsymbol{r}_{\mathrm{i}} - \boldsymbol{r}_{\mathrm{i}-1}) & \text{if } \mathrm{i} = L; \\ (\boldsymbol{r}_{\mathrm{i}+1} - \boldsymbol{r}_{\mathrm{i}-1})/2 & \text{else,} \end{cases} \tag{S9}$$

where we use the minimum-image convention for the bead position vectors $\boldsymbol{r}_{\mathrm{i}}$. We then normalize this vector to obtain the unit tangent vector $\hat{\boldsymbol{t}}_{\mathrm{i}}$. We calculate $\cos\theta_{\mathrm{nem}}$ for each bead as the scalar product of the unit tangent vectors of neighbors that lie within a sphere of size $l_{\mathrm{cut-off}}$ centered at the bead in question:

$$\cos\theta_{\mathrm{nem\ j}} = \hat{\boldsymbol{t}}_{\mathrm{j}} \cdot \hat{\boldsymbol{t}}_{\mathrm{i}}, \tag{S10}$$

where 'j' is such that $|\boldsymbol{r}_{\mathrm{j}} - \boldsymbol{r}_{\mathrm{i}}| \leq l_{\mathrm{cut-off}}$. We then average $\frac{1}{2}(3\cos^2\theta_{\mathrm{nem}} - 1)$ over all neighbours within the box to obtain the average local nematic parameter for bead 'i'.

To isolate the contribution of packing effects from the intrinsically high alignment of adjacent beads due to dsRNA persistence length, we compute a distinct inter-stem nematic order parameter. Double-stranded RNA stems are reconstructed dynamically by mapping hydrogen-bonded bead clusters and neighbors $j$ are filtered to exclude local backbone sequences ($|i-j| \leq 5$) and any beads belonging to the same reconstructed stem. Using a cutoff radius of $l_{\mathrm{cut-off}} = 4.5\,\sigma$ (where $\sigma = 0.8\,\mathrm{nm}$ is the bead diameter), the unconfined simulations yield a weak, paranematic inter-stem order of $\approx 0.1$ (Suppl. Fig. S21, left) Under confinement, this value increases to 0.12–0.13 (Suppl. Fig. S21, right), reflecting the physical alignment induced by steric compression of the rod-like stems.

While the total nematic order parameter captures the global alignment which would be in principle accessible to scattering or imaging experiments, this decoupled inter-stem parameter allows us to isolate purely topological and geometric packing constraints.

### 5 Topological characterization of RNA structures - renormalized crossing number

It is also appropriate to introduce some topological concepts relevant to pseudoknots. A graph is *reducible* if it can be decomposed into two disconnected components by cutting the backbone in one place [18, 19]. A component of a graph is *nested* if this component can be removed by cutting the backbone in two points. A *primitive* graph is one that is both irreducible and non-nested. Following Vernizzi et al. [18], we define the number of crossings in the primitive diagram of an RNA arc diagram as the *renormalized crossing number* ($N_{\mathrm{c\ renorm}}$) of the graph. We provide a table with $N_{\mathrm{c\ renorm}}$ values for our main types of simulations in Supplementary Table S4 and show arc diagrams based on simulations with stacking in which sheaves of arcs have been renormalized and arcs that have no intersections have been removed (Suppl. Fig. S13). This treatment amounts to converting an arc diagram to a sum of primitive diagrams whose genera add up to that of the original diagram. We use our own algorithm for calculating the renormalized crossing number of a base-pairing pattern and reducing its corresponding arc diagram to a sum of primitive diagrams that have $g > 0$.

We adapt the algorithms given in pseudocode in Ref. [18] for calculating the genus $g$. We test our algorithm on a subset of the PDB entries cited in Ref. [19] - 1Q81, 1QVG, 1DDY, 1NJM, 1I97, 1SJ3, 1DRZ, 1N34, from which we parse the secondary structure using the online tool RNApdbee [20]. We cross-reference the results from our implementation with the algorithm for calculating the genus by Quadrini et al. [21, 22] and find no difference. We provide additional detail on characterizing pseudoknot topology in section 1 of the SI.

### 6 Persistent pseudoknots

We provide an alternative quantification of the level of pseudoknotting in our simulations by calculating the fraction of base pairs that are involved in pseudoknots (defined as the minimum set of contacts that need to be removed from the structure to yield a pseudoknot-free arc diagram, [23]). Due to the transient nature of contacts in our simulations, we focus only on the persistent pseudoknots, which we identify by the following procedure: 1) calculating the number of frames containing each base pair seen in the simulations; 2) filtering the secondary structure for each frame, leaving only base pairs seen in at least 10% of the frames; 3) parsing the pseudoknotted base pairs via bpRNA [23]; 4) dividing the number of pseudoknotted base pairs by the overall number of base pairs in the filtered secondary structure for each frame to obtain the fraction of base pairs that form pseudoknots; 5) averaging this fraction over the last 50 simulation frames and over the 100 simulations in the ensemble. This yields 45.3±1.5% of bases in pseudoknots for confined simulations and 49.4±1.7% for unconfined simulations.

### 7 Global pseudoknot characterization

We characterize the relationship between simulated and experimental data on pseudoknots in the following way: 1) we parse all simulated frames after equilibration with bpRNA [23]; 2) for each base pair, we calculate the simulated probability that it is in a pseudoknot (i.e., the fraction of frames in which this is the case); 3) we calculate the Pearson correlation coefficient between the probabilities calculated in step 2) and the $C_{\mathrm{ij}}$ scores for the respective pairs. The resulting values, $R_{\mathrm{Pearson}} = 0.54$ for unconfined simulations and 0.46 for confined ones ($p$-value $p \approx 0$ in both cases), show that our simulations correlate with the experiments. As another way of globally characterizing the pseudoknot distributions, we provide histograms of the pseudoknot lengths for both types of simulations in Suppl. Fig. S16.

### 8 Stem length distribution analysis

We use bpRNA [23] to characterize the distribution of stem lengths in frames 50-99 of each simulation in an ensemble.

### 9 Predicted gyration radii for unconfined conditions

Our simulations with proximity ligation data input predict a more compact RNA under no confinement than those with input from RNAfold: the average gyration radius from 3D COMRADES with stacking is 38.6 $\sigma$; from 3D Vienna it is 43.9 $\sigma$.

### 10 List of supplementary movies

Movies illustrate the dynamics of the viral RNA with and without confinement. Each bead represents a nucleotide, with beads coloured according to their index starting from blue and ending with red. Cyan cylinders represent base pairs.

1. Unconfined 3D COMRADES simulation without stacking
2. Confined 3D COMRADES simulation without stacking
3. Unconfined 3D COMRADES simulation with stacking with no circularization
4. Unconfined 3D COMRADES simulation with stacking and circularization
5. Confined 3D COMRADES simulation with stacking
6. Confined 3D COMRADES simulation with stacking, focus on circularization
7. Confined 3D COMRADES simulation with stacking, electrostatics and an SBCG representation of the capsid

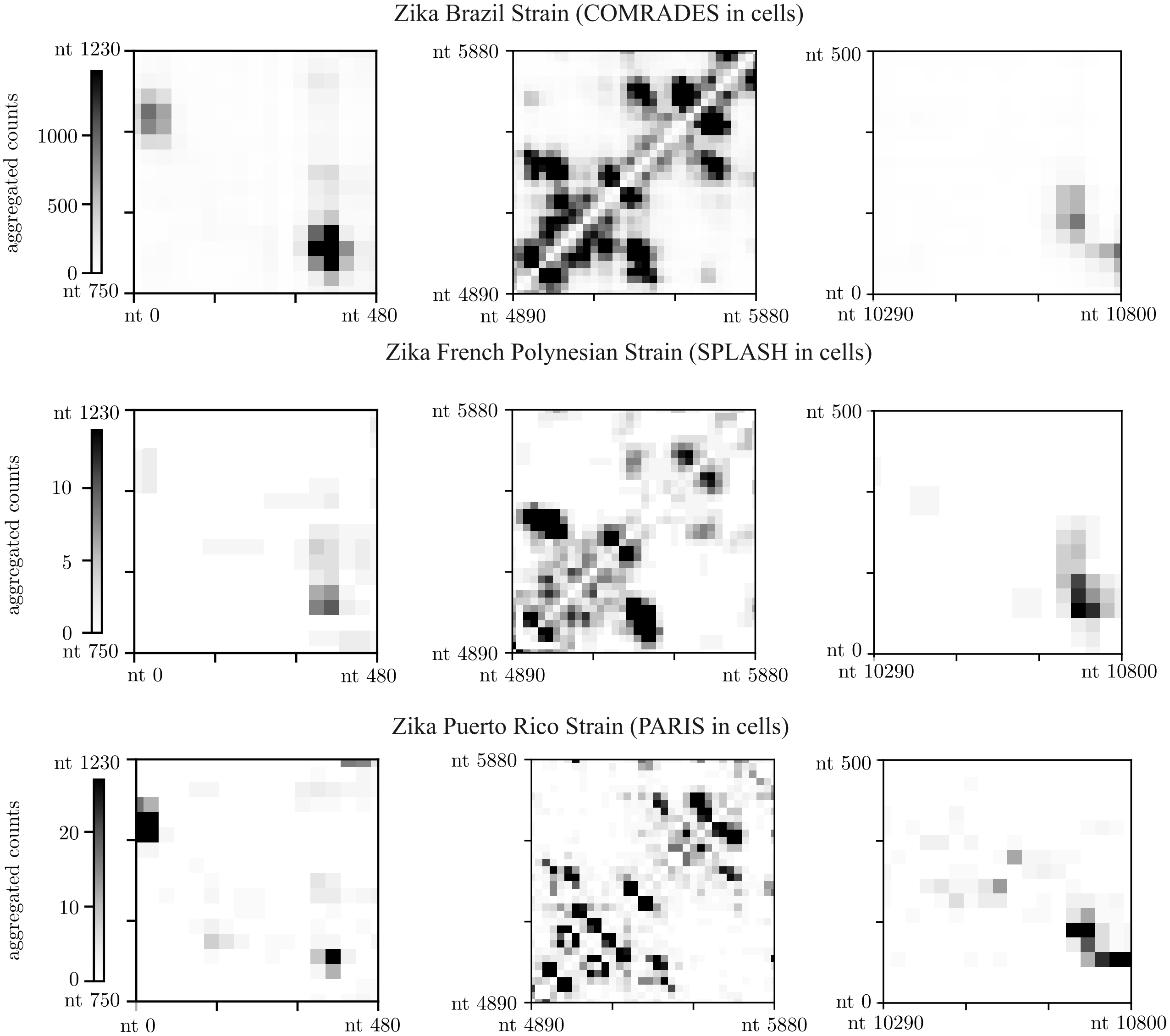


Figure S1: COMRADES, SPLASH and PARIS find consistent individual interactions for experiments conducted in cells. COMRADES data for the Brazil strain KX197192.1 [6] (top row), SPLASH data for the French Polynesia strain KJ776791.2 [8] (middle row), PARIS data for the Puerto Rico strain KX377337.1 [9] (bottom row). Regions chosen to illustrate interactions in the 500x500 nt and 1000x1000 nt windows containing 1) nt $\sim 1$ and $\sim 1000$ (leftmost column) highlighted in [9], 2) the circularization interaction (rightmost column, see also Fig. 4 in the main text) and 3) the interactions involving nt $\sim L/2 \sim 5900$, with $L$ being the length of the viral genome (centre column). Contact maps are symmetrized to show aggregated counts corresponding to signal from chimeric reads of both orientations and coarse-grained into bins of length 30 nucleotides. Contacts with aggregated counts equal to or above the 99th percentile for the specific dataset shown in black. The contact maps for the whole RNA are shown in Suppl. Fig. S4 below.

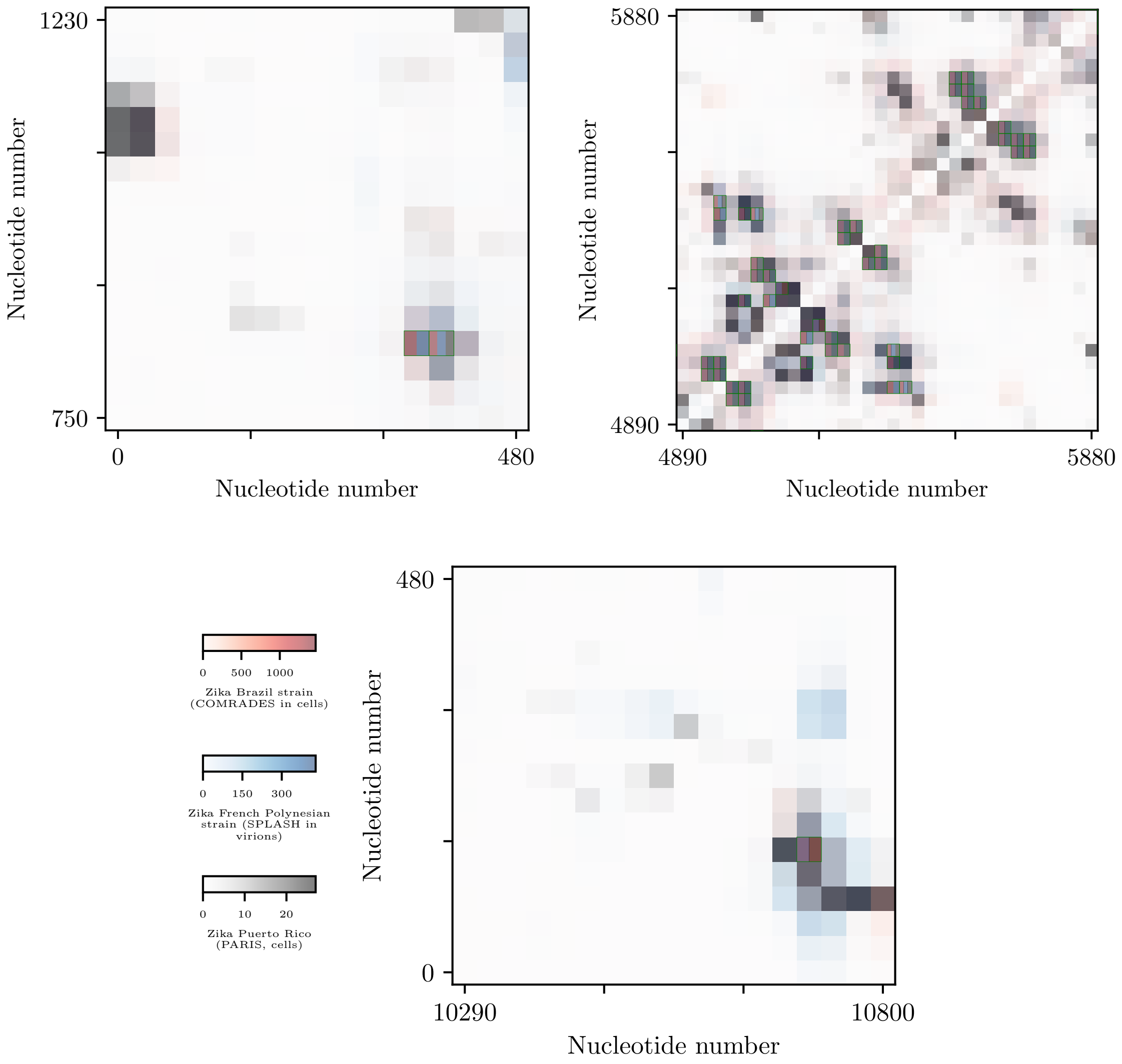


Figure S2: COMRADES, SPLASH and PARIS find consistent individual interactions for experiments conducted in cells. Contact maps as in Figure S1 above, but with each dataset plotted in a different colour: data for the Brazil strain KX197192.1 [6] (red), SPLASH data for the French Polynesia strain KJ776791.2 [8] (blue), PARIS data for the Puerto Rico strain KX377337.1 [9] (gray). Regions chosen to illustrate interactions in the 500x500 nt and 1000x1000 nt windows containing 1) nt $\sim 1$ and $\sim 1000$ (top left) highlighted in [9], 2) the circularization interaction (bottom left) and 3) the interactions involving nt $\sim L/2 \sim 5900$, with $L$ being the length of the viral genome (top right), as well as the whole RNA (next page). Contact maps are symmetrized to show aggregated counts corresponding to signal from chimeric reads of both orientations and coarse-grained into bins of length 30 nucleotides. The color scales run from zero to the 99th percentile for the specific dataset; all three layers are transparent to aid visibillity. Where the signal from two or all three datasets is equal to or greater than the respective 99th percentile, the corresponding pixel is split into two or three sections of equal area and outlined in green.

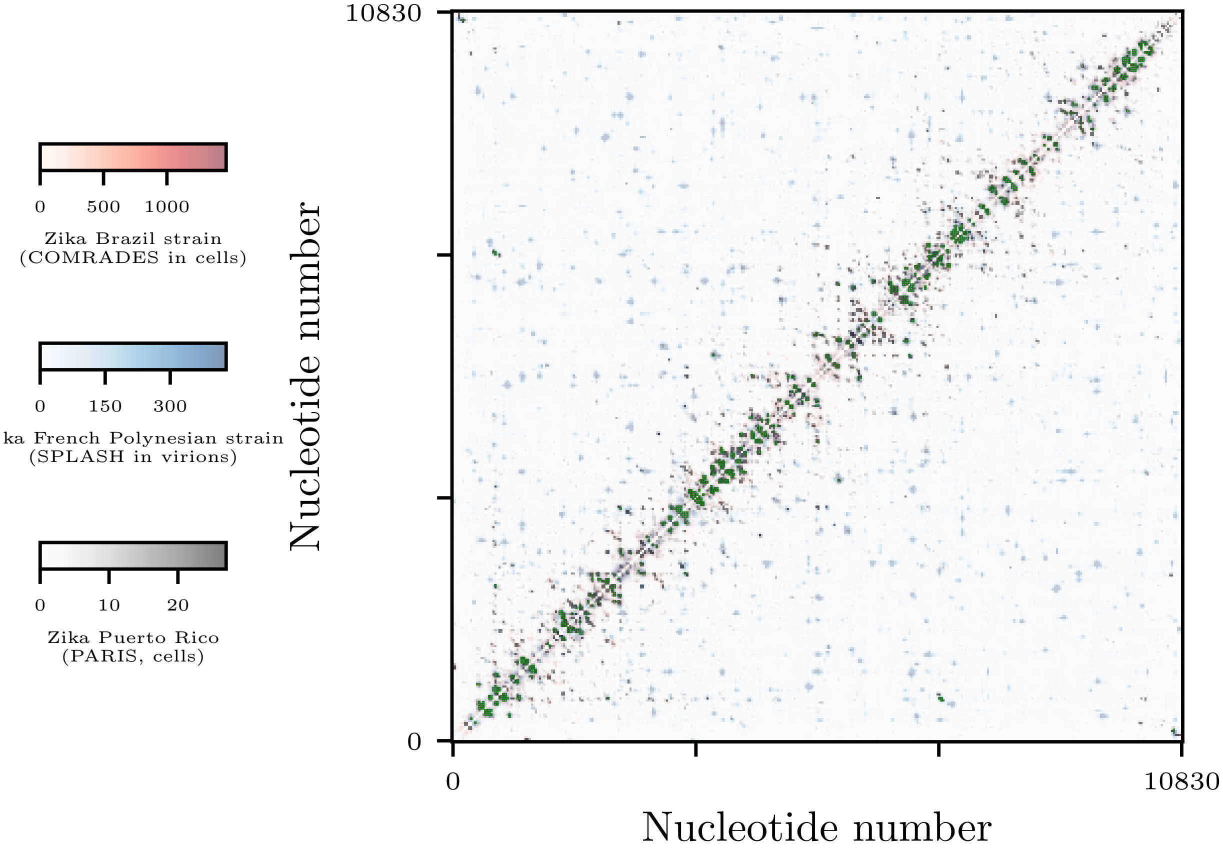
0 500 1000
Zika Brazil strain
(COMRADES in cells)
0 150 300
ka French Polynesian strain
(SPLASH in virions)
0 10 20
Zika Puerto Rico
(PARIS, cells)
10830
0
0
10830
Nucleotide number
Nucleotide number

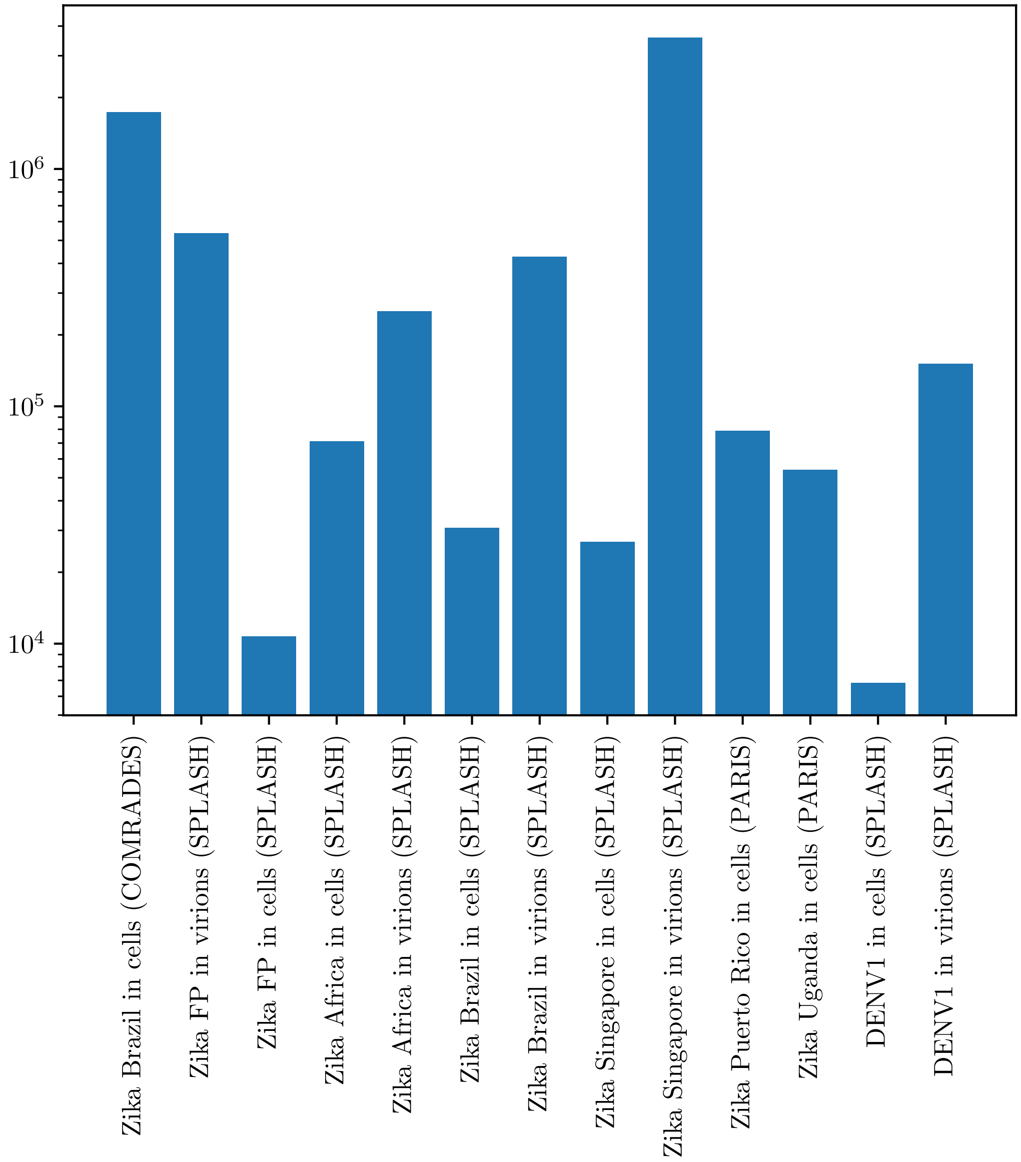


Figure S3: Number of unique chimeras that map onto the reference viral genome for all sets of experiments discussed above. Note the relatively low coverage in the SPLASH experiments with cells and the PARIS experiments.

Figure S4: *(Following pages:)* Contact maps and contact probability vs. distance plots for various type of experiments with DENV1 and various strains of the Zika virus. As in the main text, contact maps are symmetrized to show aggregated counts corresponding to signal from chimeric reads of both orientations and coarse-grained into bins of length 30 nucleotides. Contacts with aggregated counts equal to or above the 99th percentile for the specific dataset are shown in black.

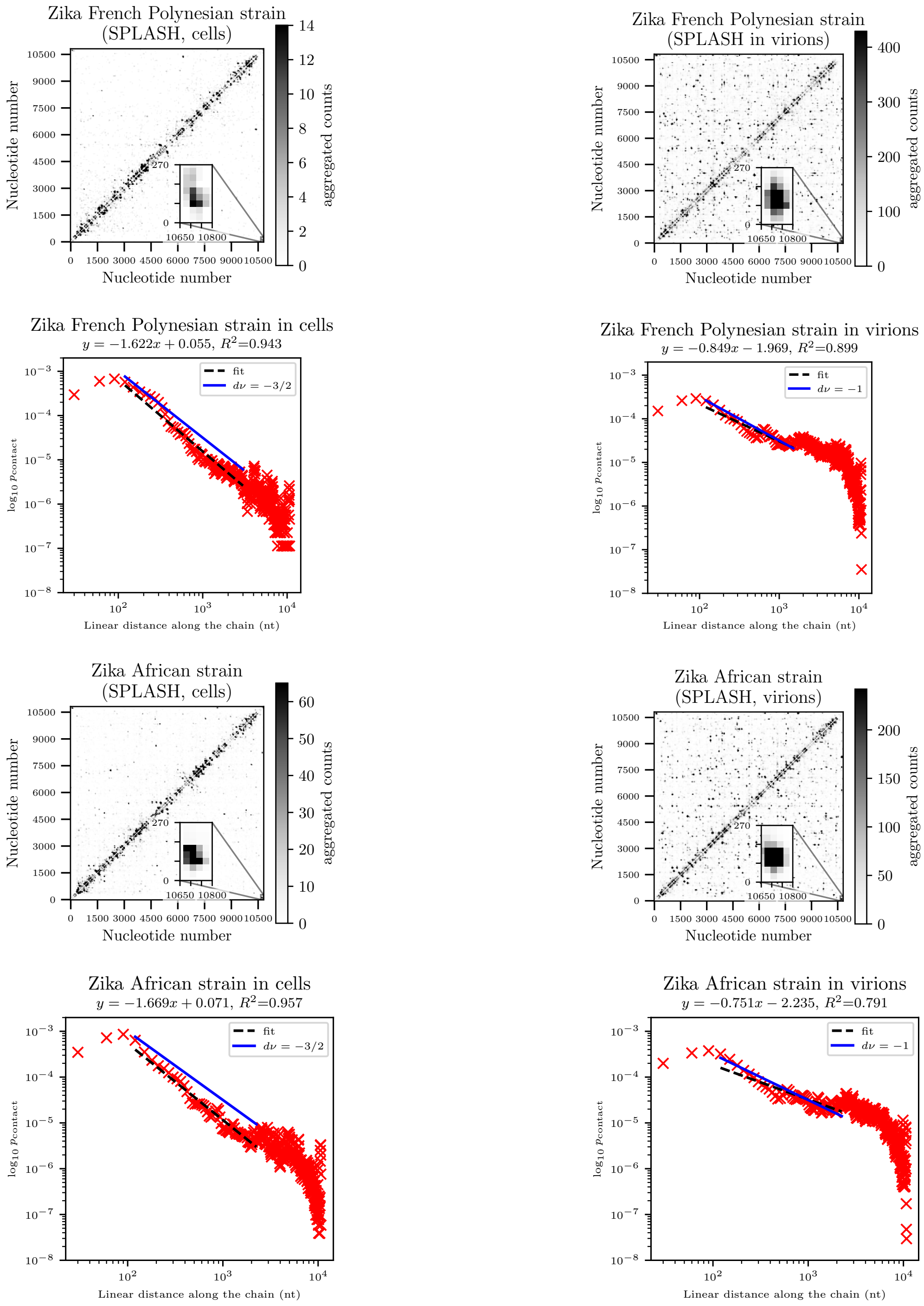
Zika French Polynesian strain (SPLASH, cells)
Nucleotide number
aggregated counts
Zika French Polynesian strain (SPLASH in virions)
Zika French Polynesian strain in cells
$y = -1.622x + 0.055$, $R^2$=0.943
fit
$d\nu = -3/2$
$\log_{10} p_{\mathrm{contact}}$
Linear distance along the chain (nt)
Zika French Polynesian strain in virions
$y = -0.849x - 1.969$, $R^2$=0.899
$d\nu = -1$
Zika African strain (SPLASH, cells)
Zika African strain (SPLASH, virions)
Zika African strain in cells
$y = -1.669x + 0.071$, $R^2$=0.957
Zika African strain in virions
$y = -0.751x - 2.235$, $R^2$=0.791

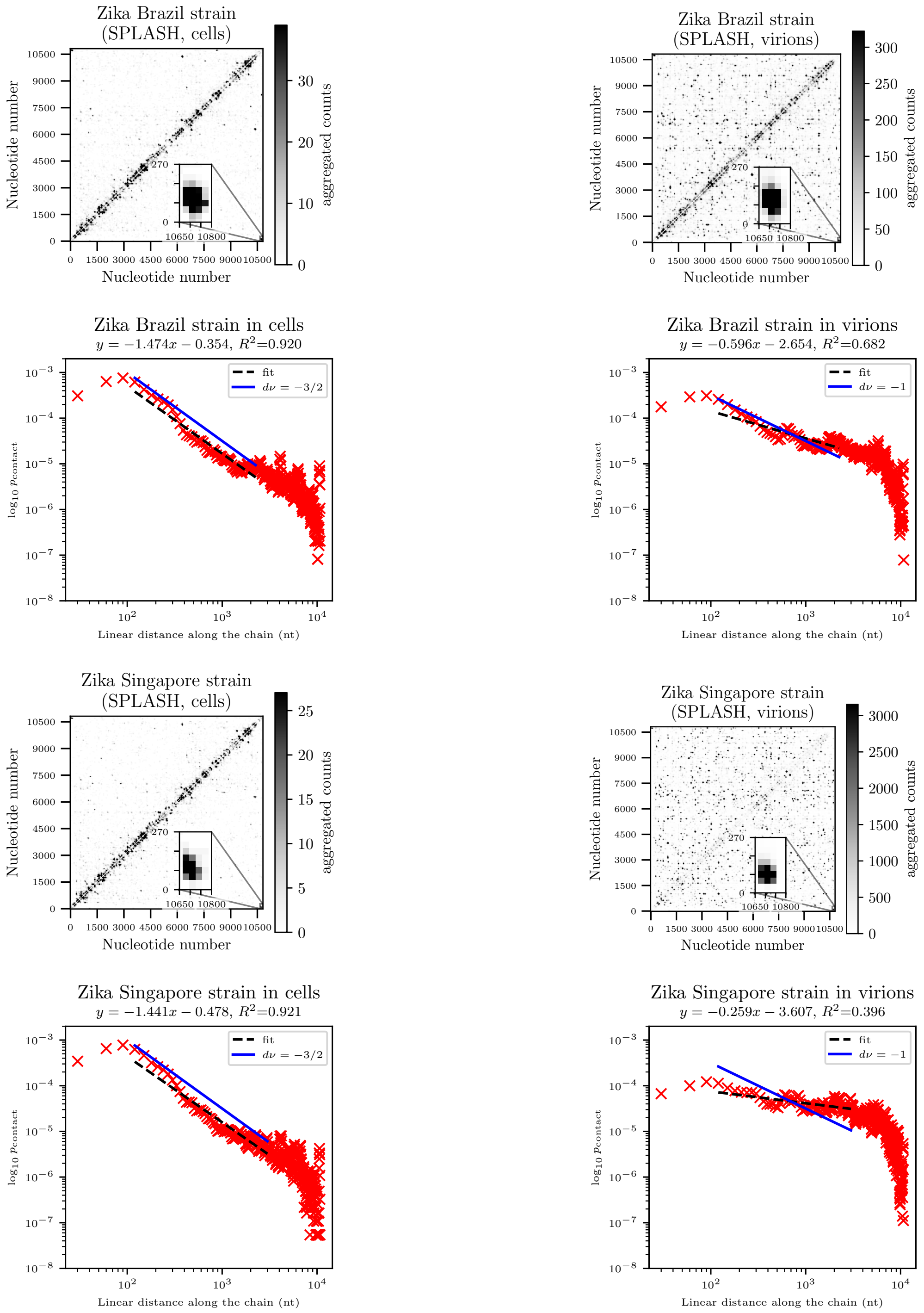
Zika Brazil strain (SPLASH, cells)
Nucleotide number
aggregated counts
Zika Brazil strain (SPLASH, virions)
Zika Brazil strain in cells
$y = -1.474x - 0.354$, $R^2$=0.920
fit
$d\nu = -3/2$
$\log_{10} p_{\mathrm{contact}}$
Linear distance along the chain (nt)
Zika Brazil strain in virions
$y = -0.596x - 2.654$, $R^2$=0.682
$d\nu = -1$
Zika Singapore strain (SPLASH, cells)
Zika Singapore strain (SPLASH, virions)
Zika Singapore strain in cells
$y = -1.441x - 0.478$, $R^2$=0.921
Zika Singapore strain in virions
$y = -0.259x - 3.607$, $R^2$=0.396

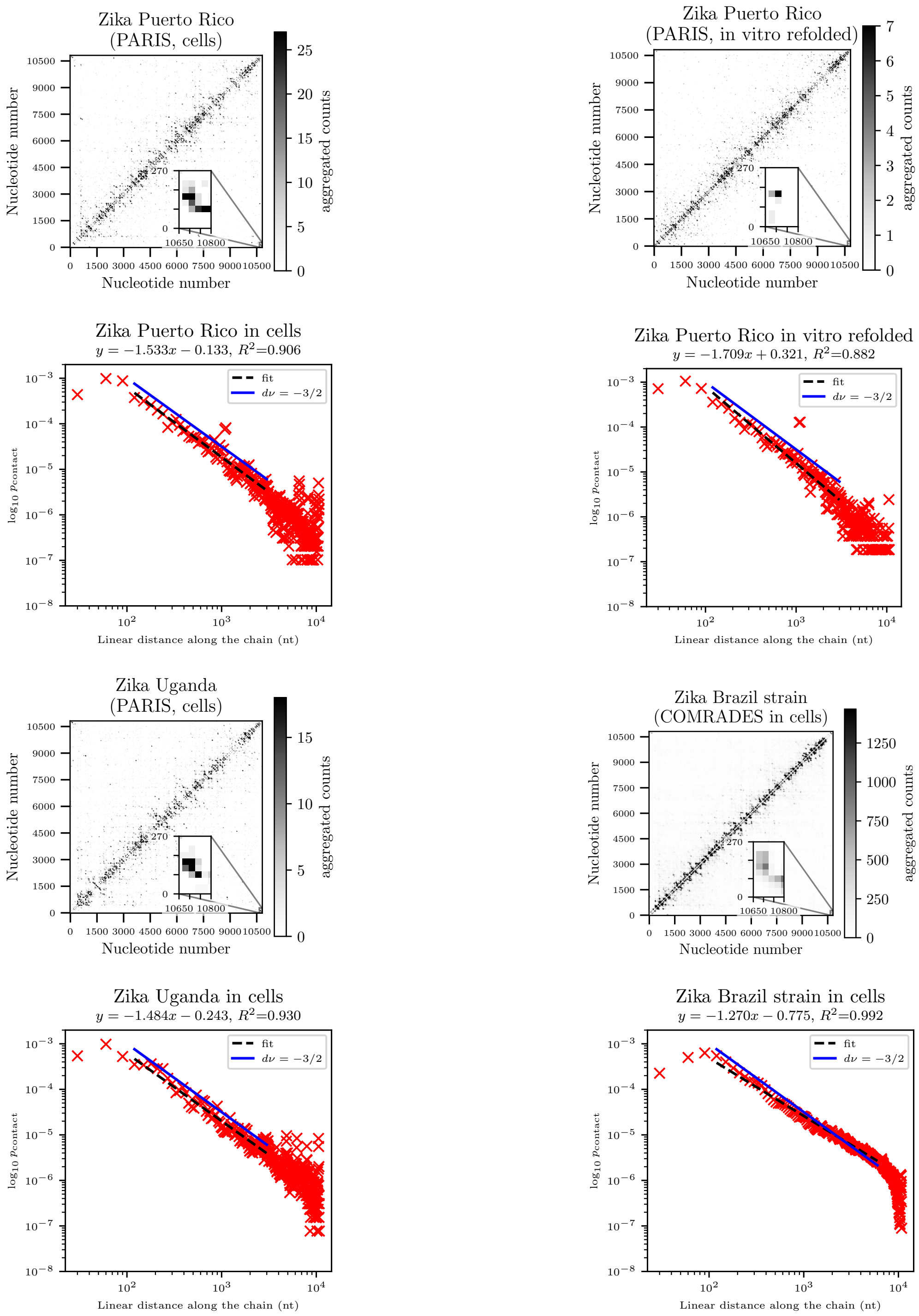
Zika Puerto Rico
(PARIS, cells)
Nucleotide number
aggregated counts
Zika Puerto Rico
(PARIS, in vitro refolded)
Nucleotide number
aggregated counts
Zika Puerto Rico in cells
$y = -1.533x - 0.133,\ R^2$=0.906
fit
$d\nu = -3/2$
$\log_{10} p_{\text{contact}}$
Linear distance along the chain (nt)
Zika Puerto Rico in vitro refolded
$y = -1.709x + 0.321,\ R^2$=0.882
fit
$d\nu = -3/2$
$\log_{10} p_{\text{contact}}$
Linear distance along the chain (nt)
Zika Uganda
(PARIS, cells)
Nucleotide number
aggregated counts
Zika Brazil strain
(COMRADES in cells)
Nucleotide number
aggregated counts
Zika Uganda in cells
$y = -1.484x - 0.243,\ R^2$=0.930
fit
$d\nu = -3/2$
$\log_{10} p_{\text{contact}}$
Linear distance along the chain (nt)
Zika Brazil strain in cells
$y = -1.270x - 0.775,\ R^2$=0.992
fit
$d\nu = -3/2$
$\log_{10} p_{\text{contact}}$
Linear distance along the chain (nt)

Figure S5: *(Following page:)* Interaction length distributions in various strains of Zika probed via COMRADES [6], SPLASH [8] and PARIS [9]. Every pairwise interaction is counted $C_{\mathrm{ij}}$ times to reflect the number of chimeras supporting it. $p$-values calculated from the Wilcoxon rank-sum test with alternative hypothesis that the interaction length is greater in datasets obtained from experiments with virions. Boxes extend from the first quartile to the third quartile of the distribution, and the median is indicated by a line. Whiskers extend from the box to the farthest data point that is within 1.5x the inter-quartile range from the box, and fliers indicate points that are farther from the median than the ends of the whiskers.

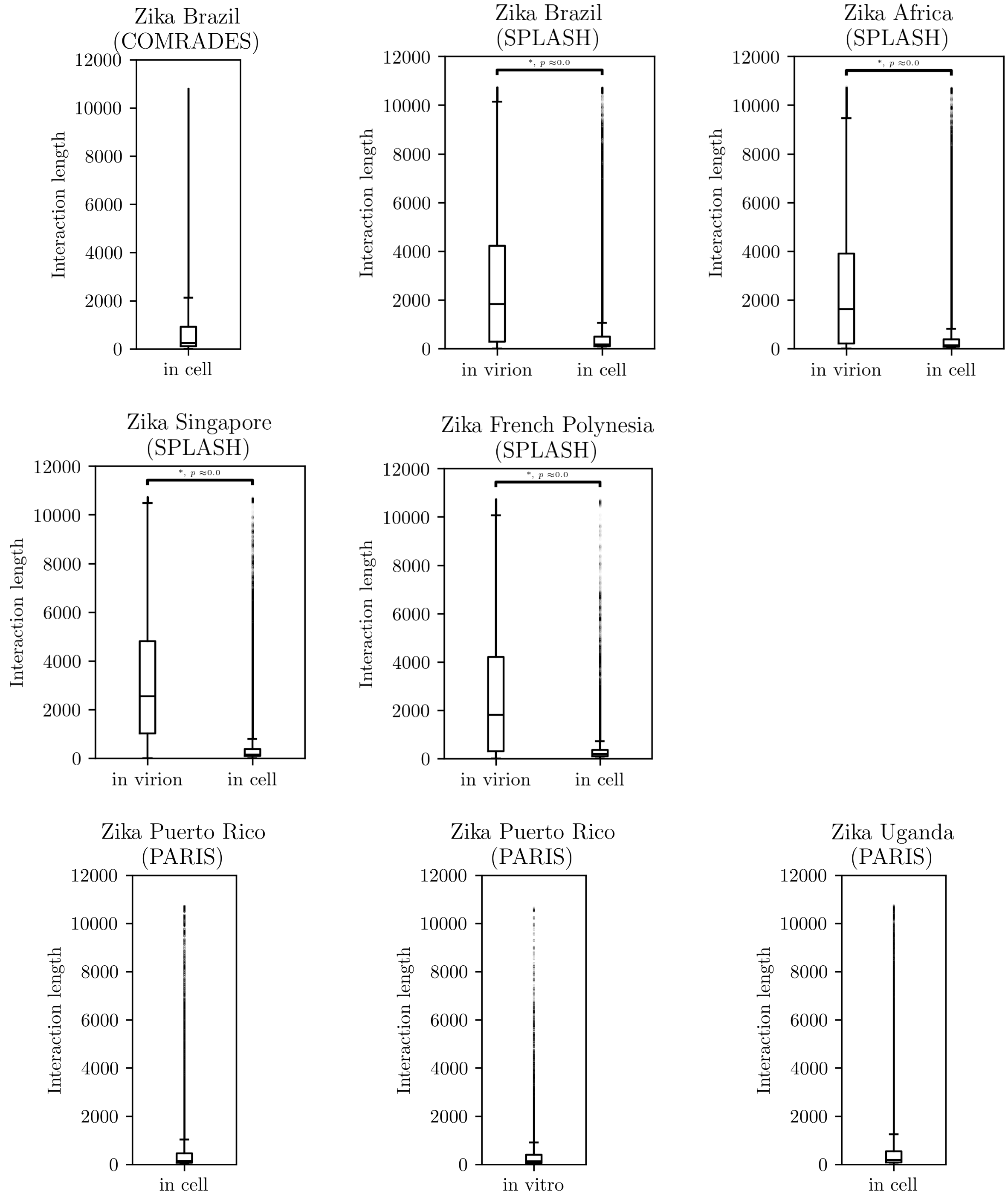

Zika Brazil
(COMRADES)
Interaction length
in cell
Zika Brazil
(SPLASH)
Interaction length
in virion
in cell
Zika Africa
(SPLASH)
Interaction length
in virion
in cell
Zika Singapore
(SPLASH)
Interaction length
in virion
in cell
Zika French Polynesia
(SPLASH)
Interaction length
in virion
in cell
Zika Puerto Rico
(PARIS)
Interaction length
in cell
Zika Puerto Rico
(PARIS)
Interaction length
in vitro
Zika Uganda
(PARIS)
Interaction length
in cell

Figure S6: Interaction length distributions in various strains of Zika probed via COMRADES [6], SPLASH [8] and PARIS [9]. Rare interactions ($C_{\text{ij}} \leq 5$) are filtered out.

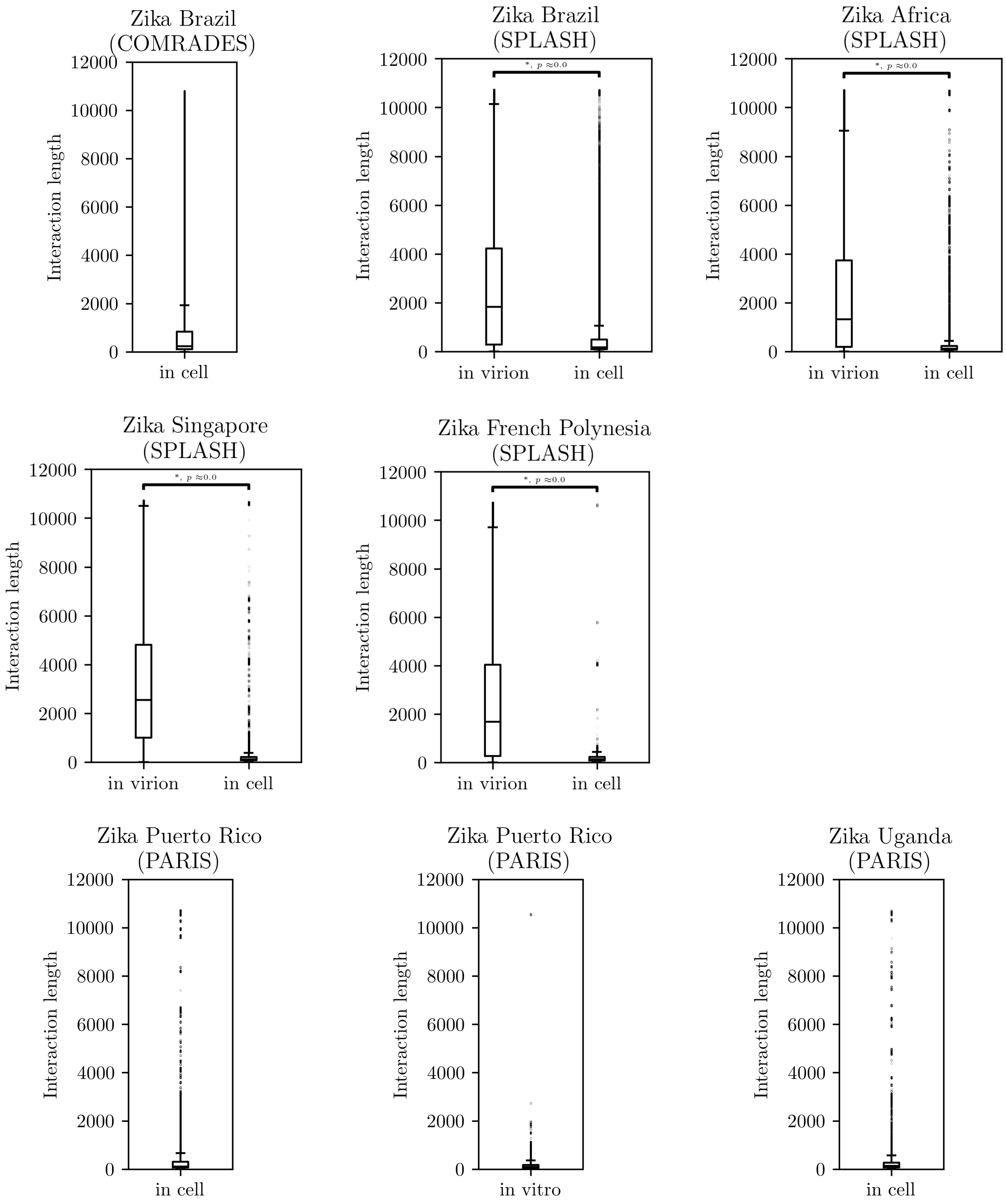

| Dataset | bin size - 30 nt | | | bin size - 10 nt | | |
|---|---|---|---|---|---|---|
| | $d\nu$ | $R^2$ | range (nt) | $d\nu$ | $R^2$ | range (nt) |
| LinearPartition, KX197192.1 | 1.666 | 0.677 | 30-1350 | 1.53 | 0.863 | 10-450 |
| RNAfold, KX197192.1 | 1.291 | 0.691 | 30-1350 | 1.803 | 0.873 | 20-550 |
| UNAFold, KX197192.1 | 2.010 | 0.556 | 30-1350 | 1.329 | 0.855 | 10-450 |
| LinearPartition, KJ776791.2 | 2.228 | 0.687 | 30-1350 | 1.475 | 0.829 | 10-450 |
| LinearPartition, KJ776791.2 with SHAPE-MaP input | 2.010 | 0.432 | 30-1500 | 2.244 | 0.444 | 0-1000 |
| RNAfold, KJ776791.2 | 1.252 | 0.642 | 30-1650 | 1.618 | 0.877 | 10-550 |
| RNAfold, KJ776791.2 with SHAPE-MaP input | 1.541 | 0.636 | 30-1650 | 1.711 | 0.800 | 10-550 |
| UNAFold, KJ776791.2 | 1.783 | 0.609 | 30-1500 | 1.836 | 0.561 | 10-1500 |

Table S1: Critical exponents determined from fits of $p_{\text{contact}}$ vs. genomic distance for secondary structure prediction algorithms applied to the Zika sequences used for in vivo experiments (KX197192.1) and ones in virions (KJ776791.2).

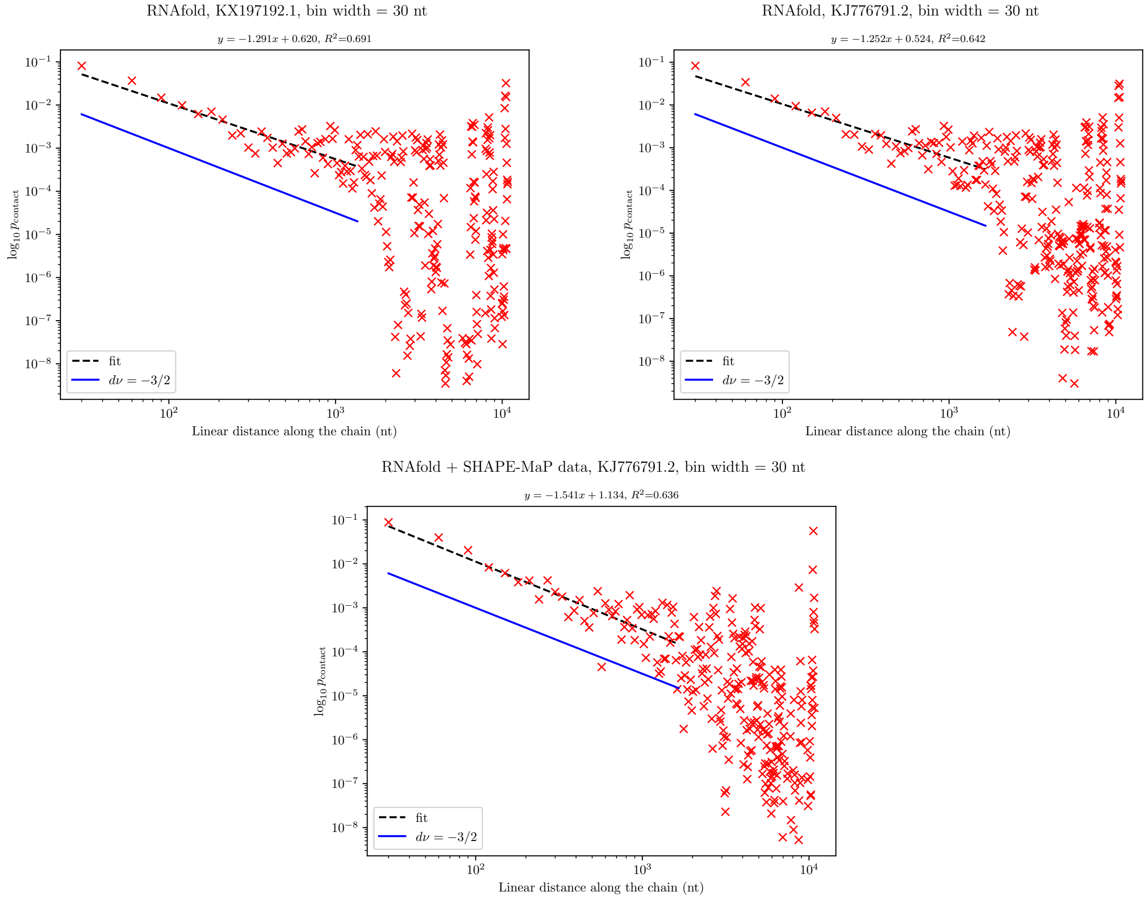


Figure S7: The ViennaRNA RNAfold program predicts similar slopes $-d\nu \sim -1.5$ for all conditions in the log-log plot of contact probability versus genomic distance, close to the expectation for an ideal chain. Ensemble base-pairing predictions for the KX197192.1 Zika sequence used in generating in vivo contact maps [6] (top left), the KJ776791.2 sequence used in experiments with virions [8], and the KJ776791.2 together with SHAPE-MaP input from Huber et al. [8]. All data is coarse-grained into bins of width 30 nucleotides.

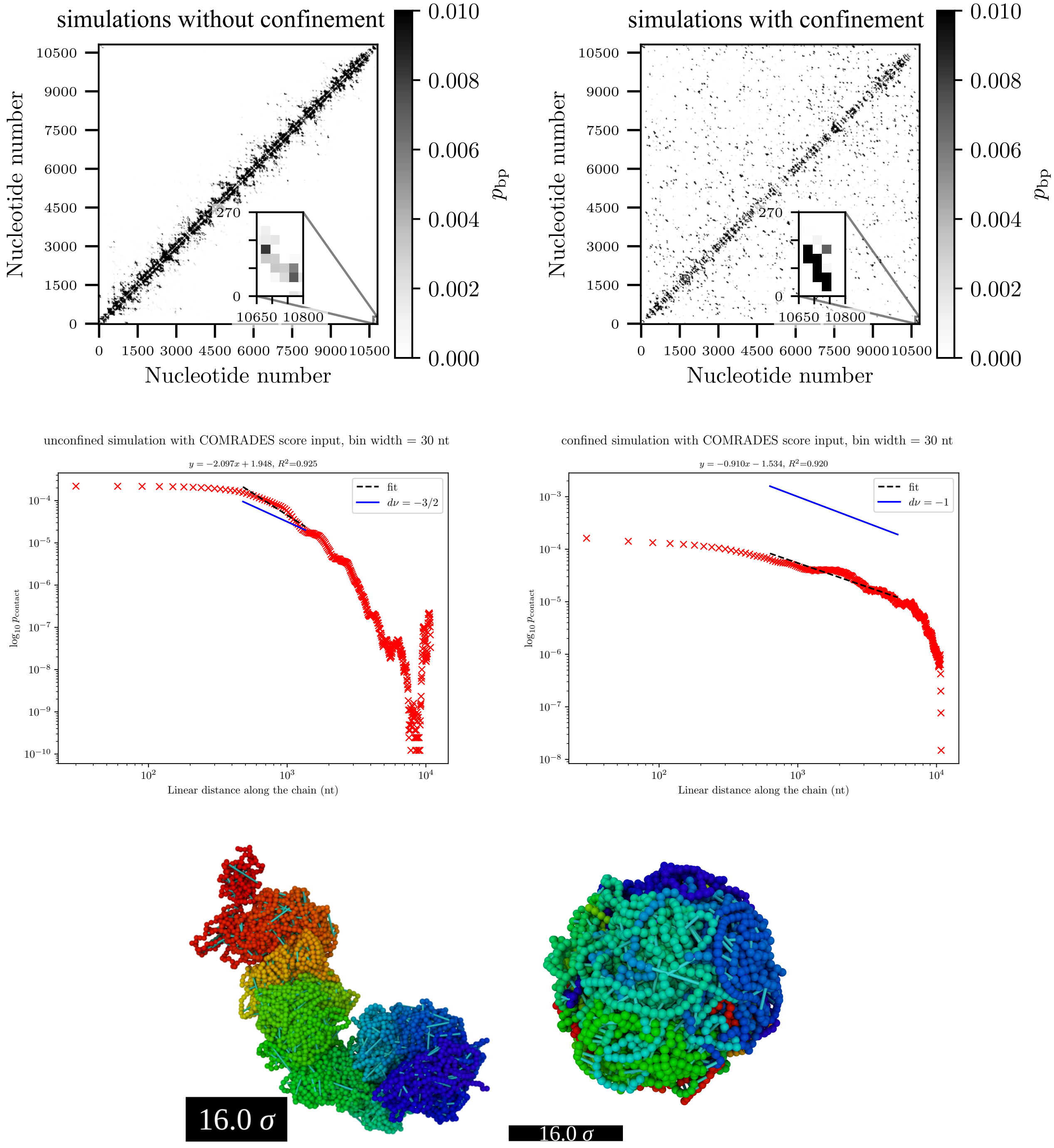
simulations without confinement
simulations with confinement
Nucleotide number
$p_{bp}$
0.010
0.008
0.006
0.004
0.002
0.000
270
10650 10800
unconfined simulation with COMRADES score input, bin width = 30 nt
$y = -2.097x + 1.948$, $R^2$=0.925
fit
$d\nu = -3/2$
confined simulation with COMRADES score input, bin width = 30 nt
$y = -0.910x - 1.534$, $R^2$=0.920
$d\nu = -1$
$\log_{10} p_{\rm contact}$
Linear distance along the chain (nt)
16.0 σ
16.0 σ

Figure S8: *(previous page)* Ensemble- and time-averaged simulated contact maps (top row) and contact probability vs genomic distance curves with a contact defined as occurring when the distance between bin centres-of-mass is less than $d_{\text{cont}} = 10\sigma$ (middle row), where $\sigma$ is the diameter of a coarse-grained bead. Simulated data is coarse-grained with a bin size of 30 nt. Snapshots of the final frame (bottom row) of a simulation without confinement (left) and with confinement (right), with the beads coloured according to their index starting from blue and ending with red (scale bar length approximately equal to the radius of the inner capsid of the Zika virus). These simulations, which do not account for base-pair stacking, correlate well with experimental data but display highly stretched base pairs (base pairs indicated in cyan) due to frequent pairing of neighbouring bases to distant regions of the molecule.

| Type of simulation | $d\nu$ | $R^2$ | range (nt) |
|---|---|---|---|
| unconfined, RNAfold input | 1.630 | 0.747 | 30-600 |
| confined, RNAfold input | 1.619 | 0.760 | 30-600 |
| unconfined, COMRADES score input, no stacking | 2.809 | 0.882 | 180-1350 |
| confined, COMRADES score input, no stacking | 0.941 | 0.685 | 120-1950 |
| unconfined, COMRADES score input and stacking | 3.255 | 0.738 | 120-1200 |
| confined, COMRADES score input and stacking | 1.585 | 0.854 | 120-1500 |

Table S2: Critical exponents determined from fits of $p_{\text{contact}}$ vs. genomic distance for various types of simulations, with contact defined through base-pairing. Data is binned in bins of size 30 nt.

| Type of simulation | bin size - 30 nt | | | bin size - 10 nt | | |
|---|---|---|---|---|---|---|
| | $d\nu$ | $R^2$ | range (nt) | $d\nu$ | $R^2$ | range (nt) |
| unconfined, no base-pairing | 1.365 | 0.986 | 30-1650 | 1.267 | 0.995 | 10-1200 |
| confined, no base-pairing | 1.132 | 0.981 | 930-7500 | 1.095 | 0.982 | 910-7500 |
| unconfined, RNAfold input | 1.382 | 0.949 | 120-4500 | 1.359 | 0.964 | 130-4500 |
| confined, RNAfold input | 0.807 | 0.978 | 150-5250 | 0.767 | 0.977 | 130-5250 |
| unconfined, COMRADES score input, no stacking | 2.097 | 0.925 | 480-1350 | 1.737 | 0.942 | 460-1350 |
| unconfined, COMRADES score input, no stacking | 0.910 | 0.920 | 630-5250 | 0.910 | 0.925 | 760-5250 |
| unconfined, COMRADES score input and stacking | 3.018 | 0.997 | 480-1800 | 2.633 | 0.994 | 460-1800 |
| confined, COMRADES score input and stacking | 0.943 | 0.990 | 480-6000 | 0.870 | 0.990 | 460-6000 |

Table S3: Critical exponents determined from fits of $p_{\text{contact}}$ vs. genomic distance for various types of simulations, with contact defined as two beads coming closer than $d_{\text{cont}} = 10\sigma$ of each other.

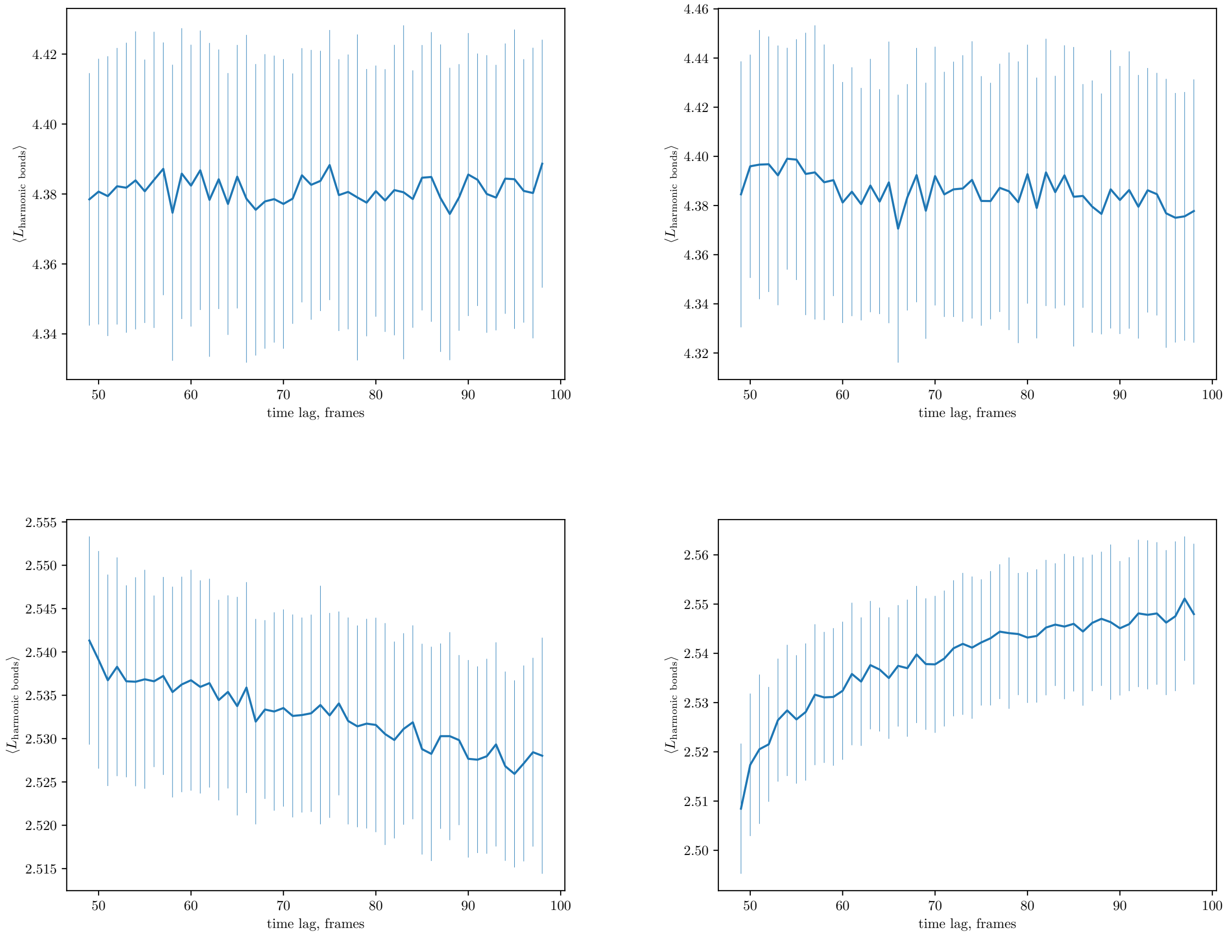


Figure S9: The average length of harmonic bonds representing base pairs is far from the equilibrium value of $2.4\sigma$ in simulations with no stacking (top row) and close to it in simulations with stacking (bottom row). The stretching of these harmonic bonds is reduced stacking and penalizing the pairing of neighbouring bases to distant regions of the molecule. Data from unconfined simulations on the left and confined ones on the right, with all ensembles using COMRADES score input and containing 100 runs. Error bars indicate standard deviation of the average length in the ensemble.

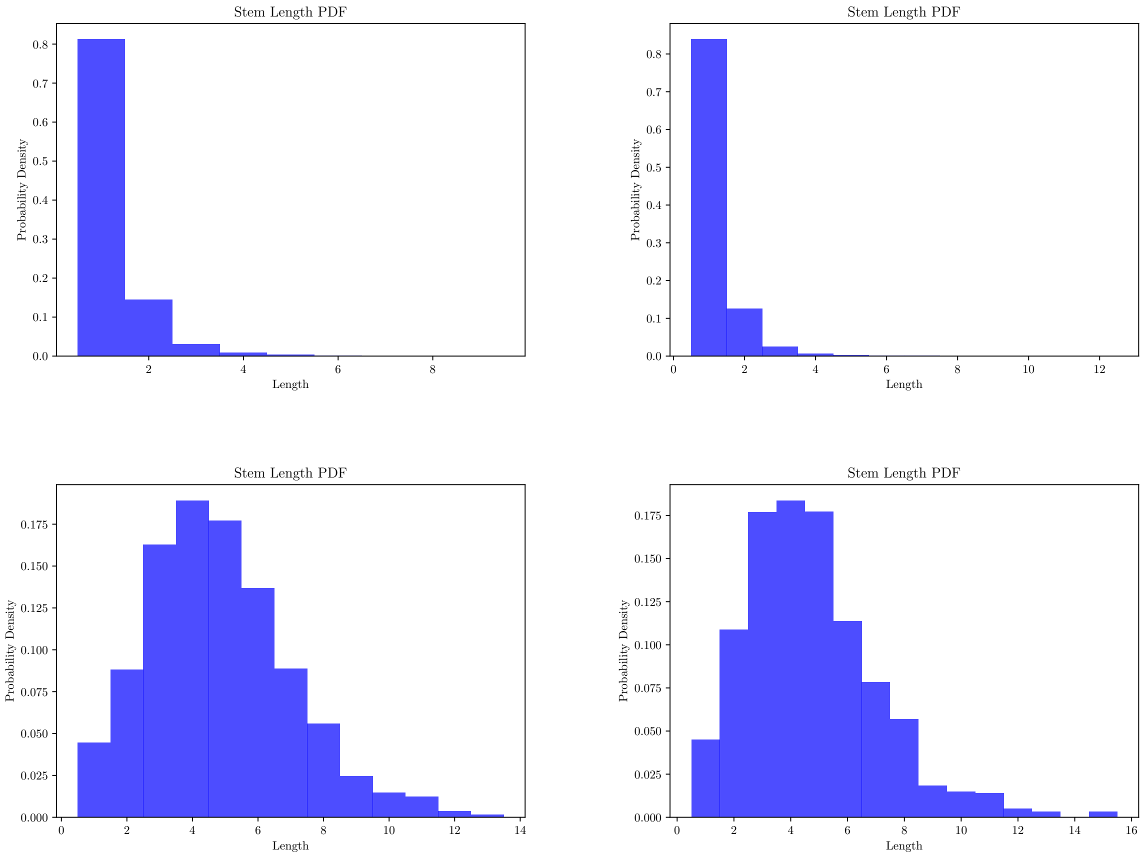


Figure S10: Including an account of stacking in the model promotes the formation of long double-stranded stems. Histograms of the stem length distributions in ensembles of unconfined (left) and confined simulations (right) with no stacking (top) and with stacking (bottom). Stems parsed with bpRNA [23].

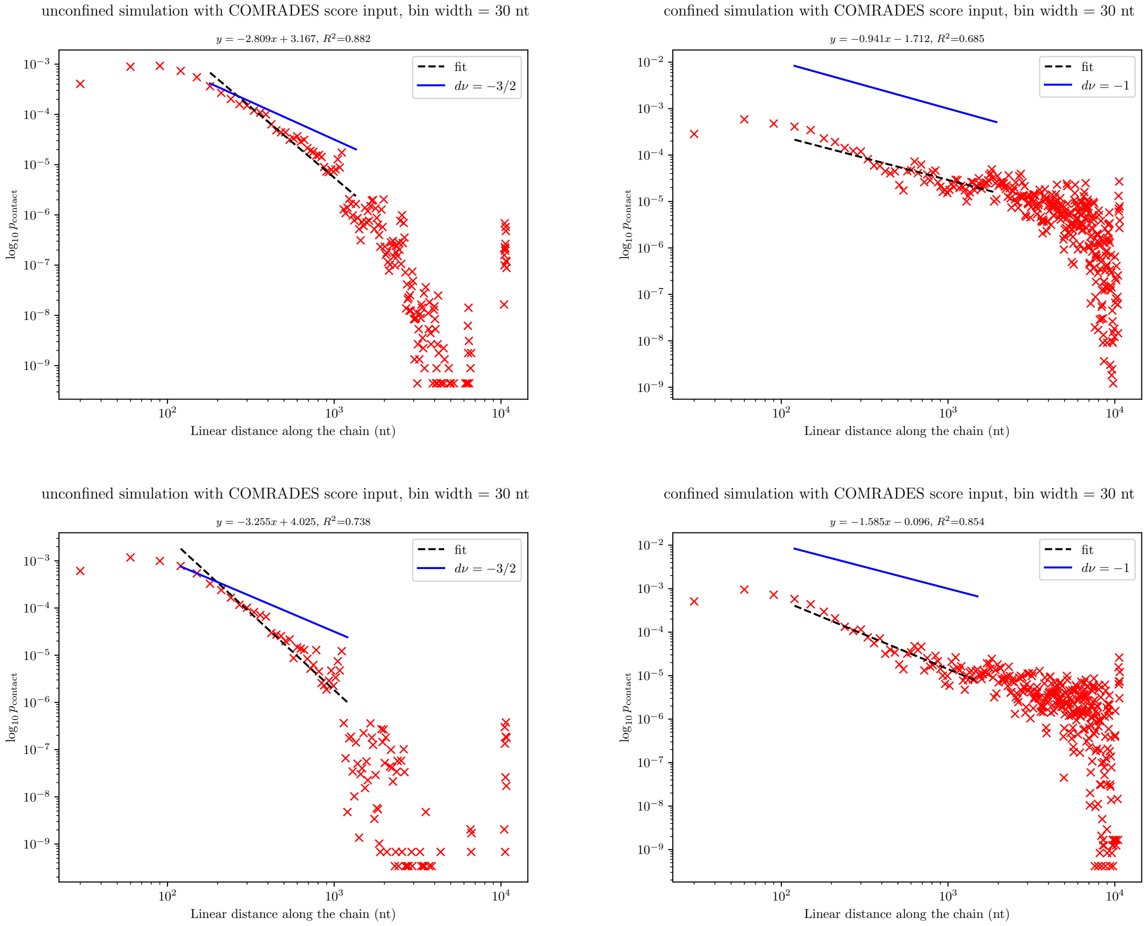


Figure S11: Simulated contact probability vs genomic distance curves for unconfined (left) and confined (right) conditions with contacts defined through base-pairing for contact maps coarse-grained with a bin width of 30 nt. Data from ensembles of simulations with (top row) and without stacking (bottom row).

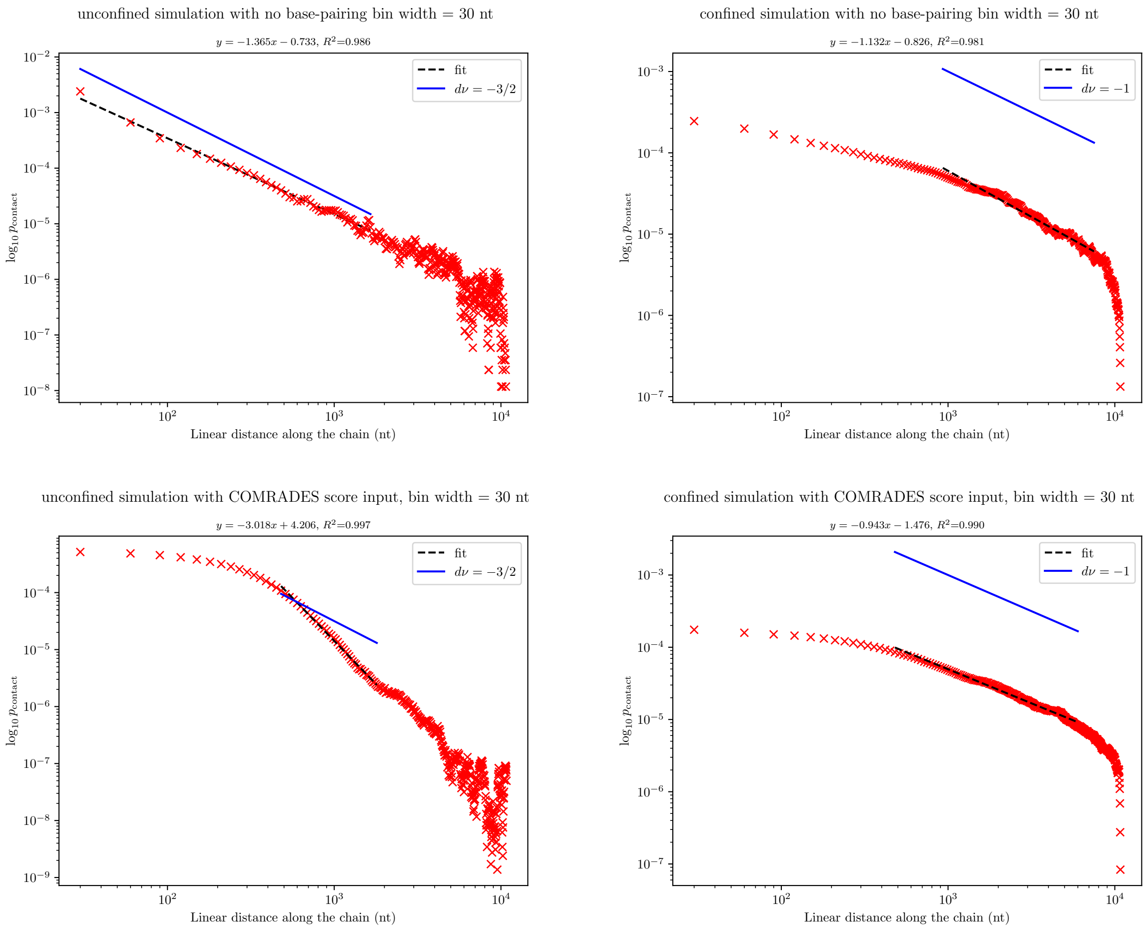


Figure S12: Contact probability vs genomic distance curves with a contact defined as occurring when the distance between bin centres-of-mass is less than $d_{\mathrm{cont}} = 10\sigma$ for simulations with and without stacking. Data from ensembles of simulations with no base-pairing (top row), and with COMRADES input and stacking (bottom row). Data is binned in bins of size 30 beads.

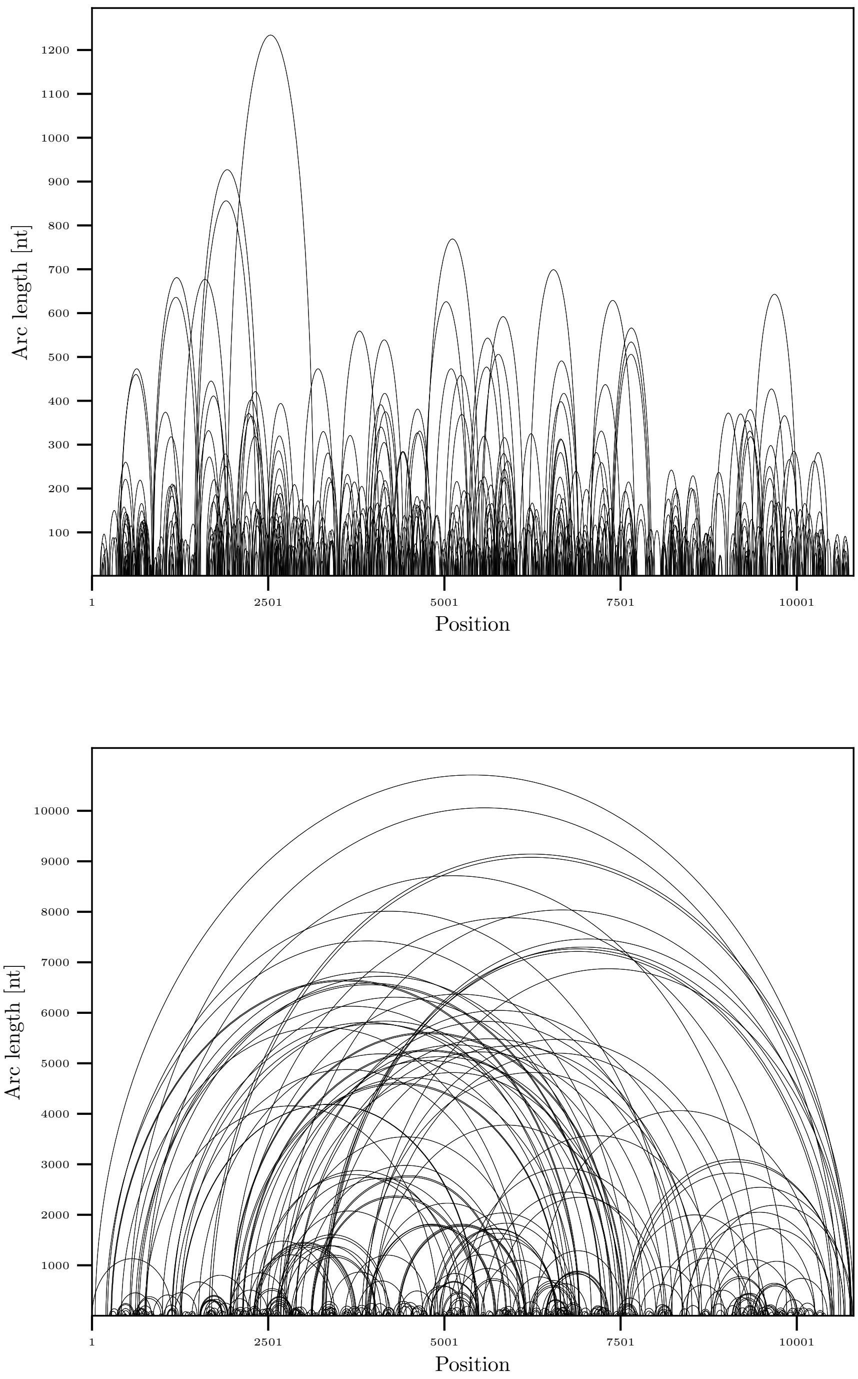


Figure S13: Arc diagrams of the structures seen in frame 99 in an unconfined (top) and confined (bottom) simulation with stacking and COMRADES score input. Sheaves of parallel arcs are shown as single renormalized arcs. Note the different vertical scale and the prominent long-range interactions in the confined case.

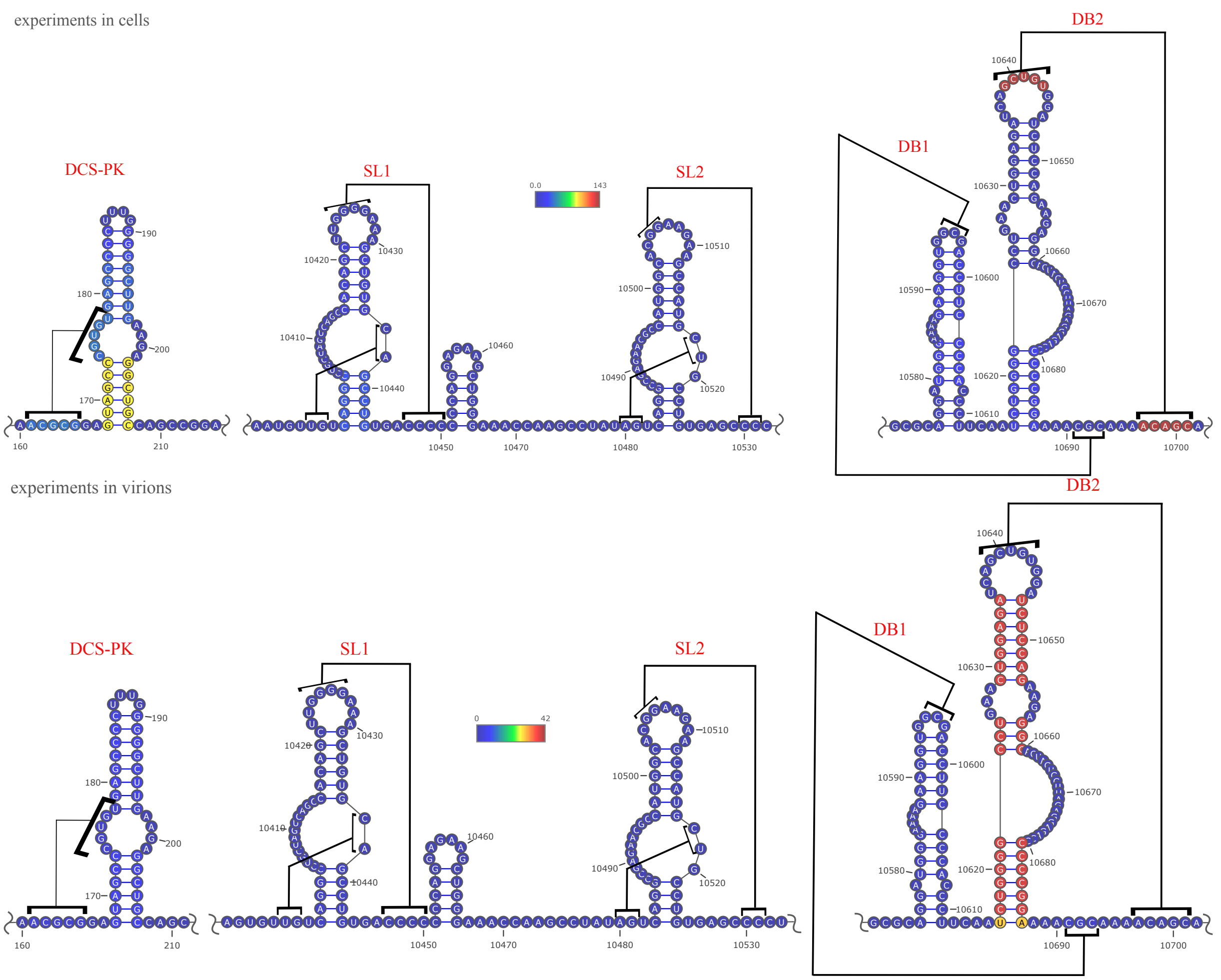


Figure S14: Proximity ligation assay data for cells in from COMRADES (top) and for virions from SPLASH (bottom) show relatively little support for well-characterized pseudoknots in the Zika virus. Pseudoknot annotations following [9], base pairs coloured according to the $C_{\mathrm{ij}}$ scores supporting them in the respective experiment. Downstream of the $5'$-cyclization sequence pseudoknot (DCS-PK), stem-loop 1 and 2 pseudoknots (SL1, SL2), and dumbbell 1 and 2 pseudoknots (DB1, DB2).

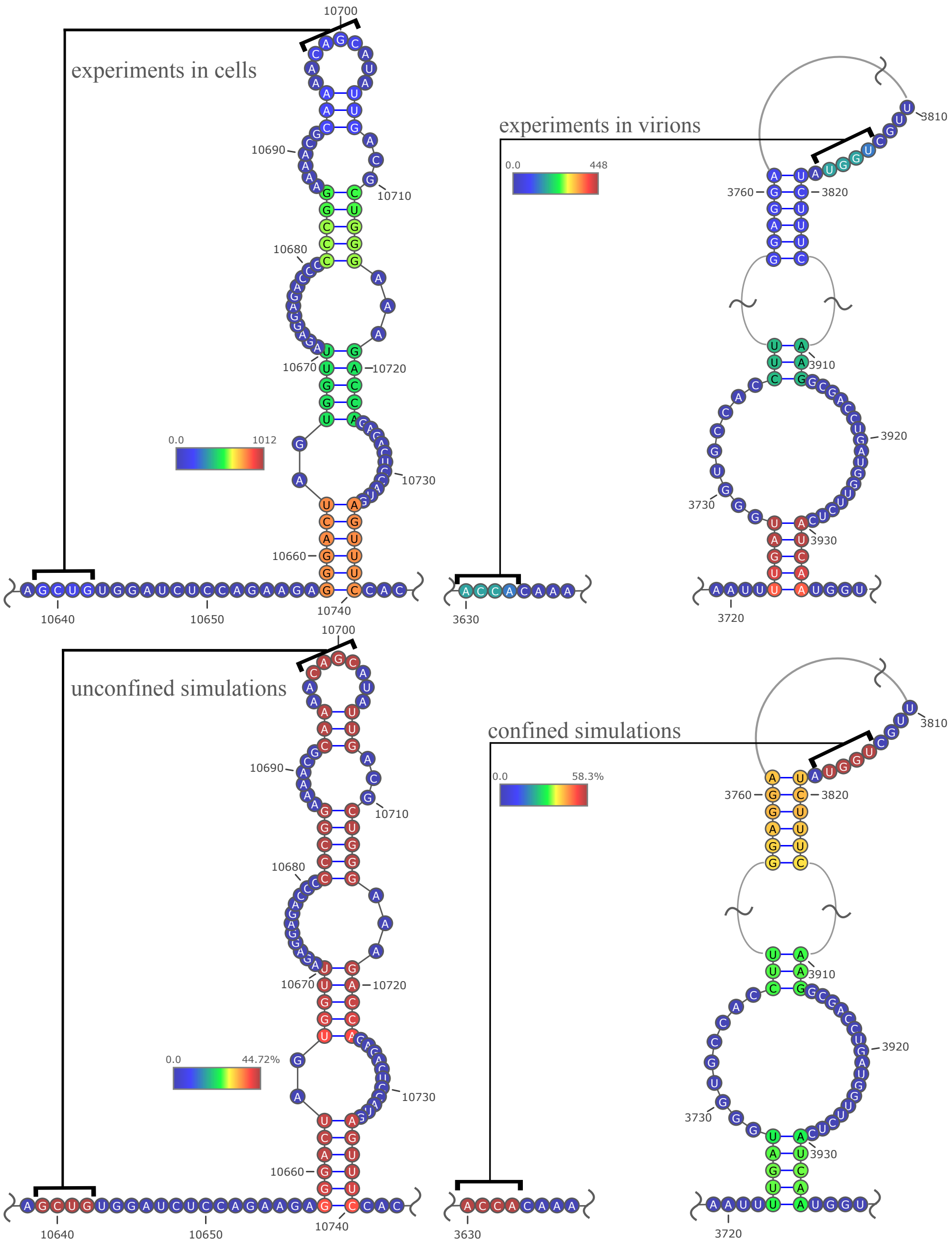


Figure S15: Potential newly identified pseudoknots in proximity ligation (top) and simulated data (bottom). These pseudoknots are supported by much greater numbers of chimeric reads in experimental data than the well-known ones shown in Suppl. Fig. S14. Base pairs coloured according to A) the $C_{\mathrm{ij}}$ scores supporting them in the respective experiment for experimental data or B) the fraction of simulation frames containing the base pair for simulated data. The example structures are chosen because they are the most frequent pseudoknots found in the respective type of simulation. For unconfined simulations, we show the contacts that are pseudoknotted with the stem consisting of bases 10639-10642 paired to 10701-10698 and occur in at least 30% of simulation frames. For confined simulations, we display the equivalent for the stem involving bases 3630-3633 and 3817-3814.

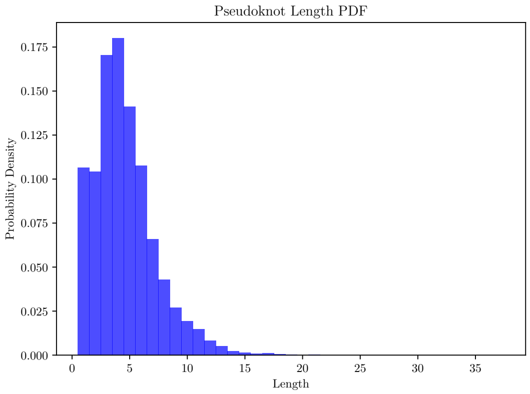


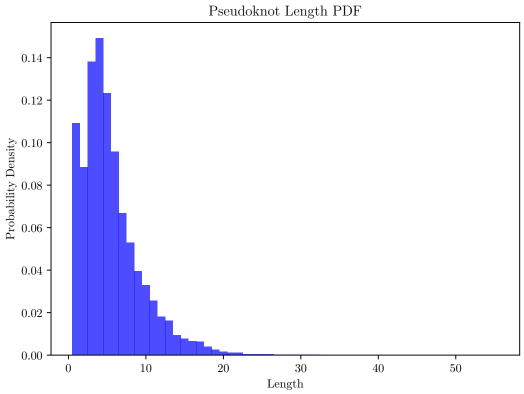


Figure S16: Histograms of the pseudoknot length distributions in ensembles of unconfined (left) and confined simulations (right) with stacking. Pseudoknots parsed with bpRNA [23].

| Dataset | $\langle N_{\text{c renorm}} \rangle$ | $\text{SD}_{\langle N_{\text{c renorm}} \rangle}$ |
|---|---|---|
| no confinement, RNAfold input | 112.7 | 22.5 |
| unconfined 3D COMRADES, no stacking | $4.69 \times 10^4$ | $1.33 \times 10^3$ |
| unconfined 3D COMRADES with stacking | $2.45 \times 10^3$ | $2.28 \times 10^2$ |
| confinement, RNAfold input | 169.5 | 26.8 |
| confined 3D COMRADES, no stacking | $1.28 \times 10^5$ | $1.34 \times 10^4$ |
| confined 3D COMRADES with stacking | $4.40 \times 10^3$ | $5.50 \times 10^2$ |

Table S4: Ensemble-averaged number of crossings between renormalized arcs for frame 99 in various types of simulations.

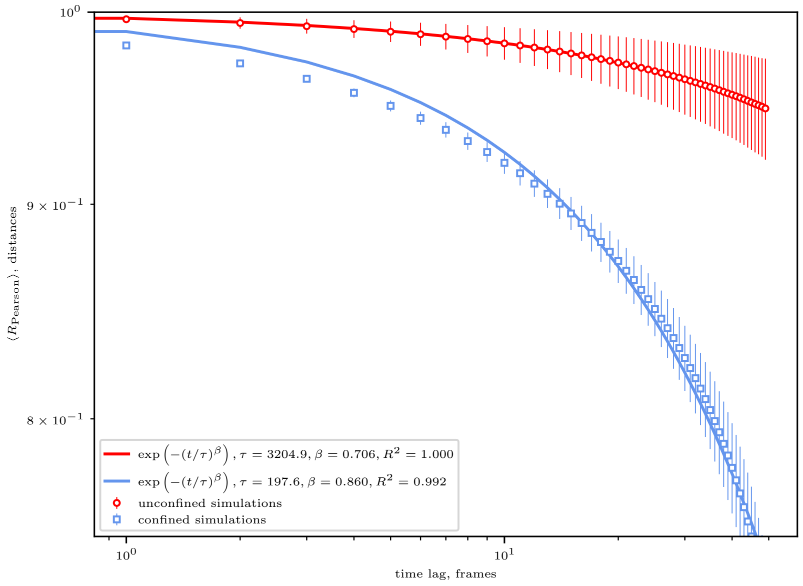


Figure S17: Interbead distances remain strongly correlated at long times and suggest glassy behaviour, similarly to base-pairing correlations (see Fig. 3 in the main text). Beads are binned into bins of size 10 nt.

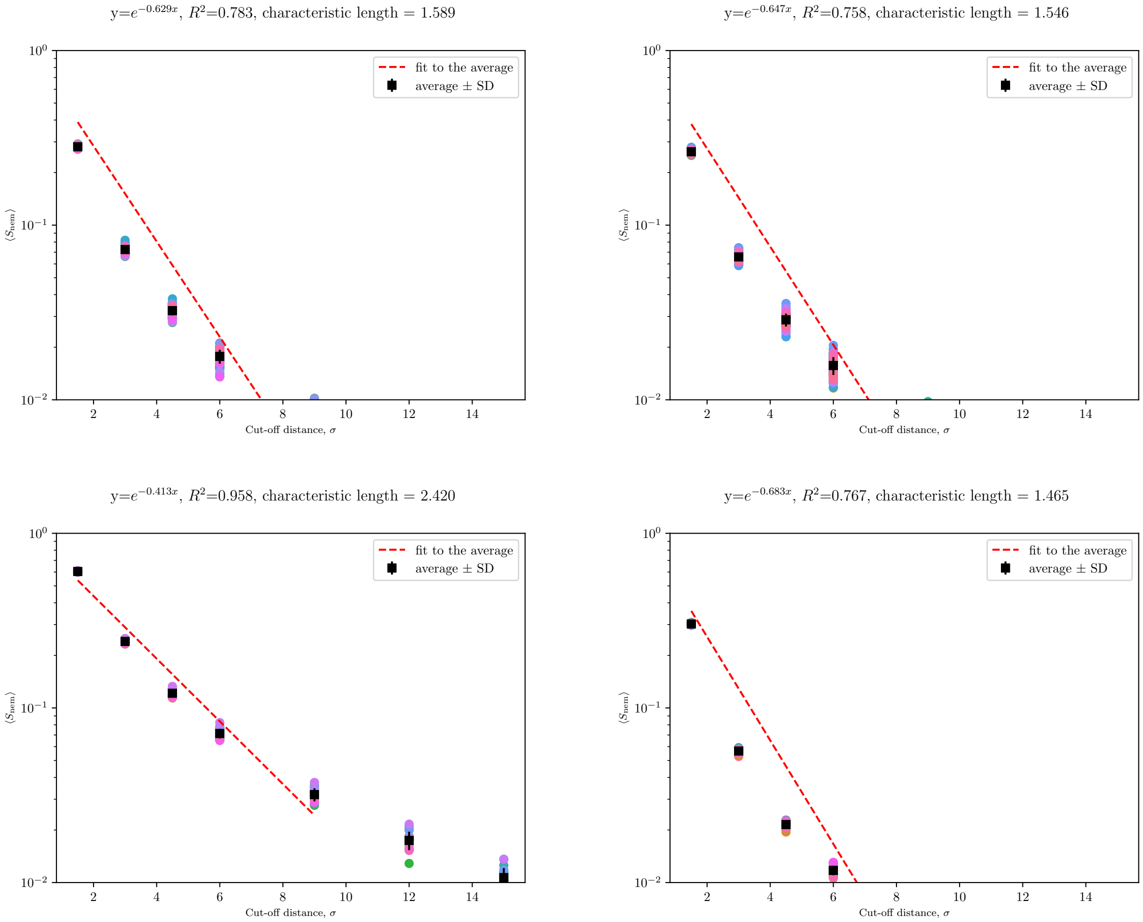


Figure S18: $\langle S_{\mathrm{nem}} \rangle$ vs cut-off distance for the last simulation frame for 1) ensembles of simulations with COMRADES score input and no stacking that are unconfined (top left) and confined (top right), and 2) ensembles of simulations with no base pairing without confinement (bottom left) and with confinement (bottom right). Each run is represented with symbols of a different colour.

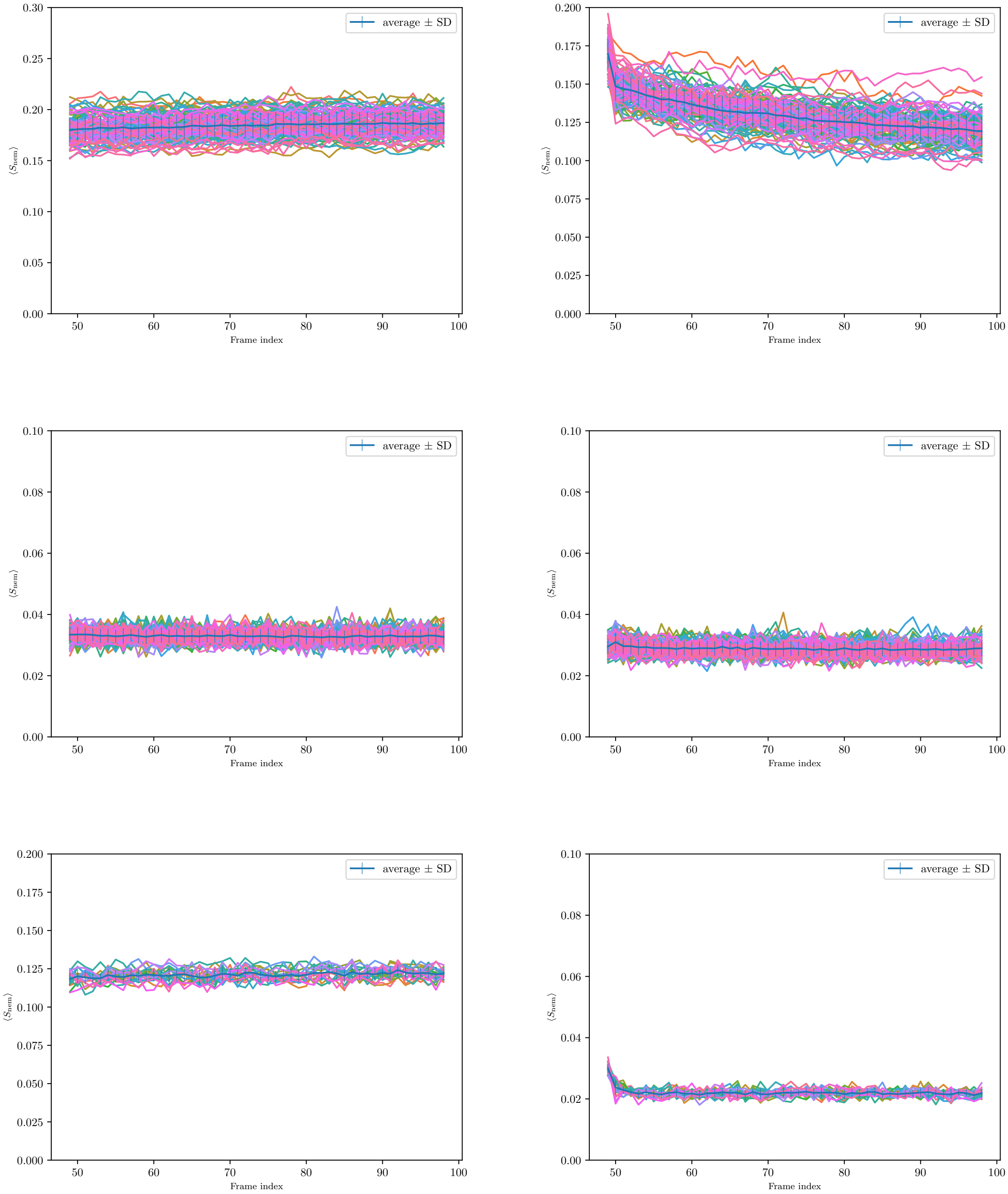


Figure S19: $\langle S_{\text{nem}}(t)\rangle$ for $l_{\text{cut-off}} = 4.5\sigma$ and ensembles of simulations 1) with COMRADES score input and stacking (top row), with COMRADES score input and no stacking (middle row), with no base-pairing (bottom row), with unconfined conditions on the left and confined - on the right.

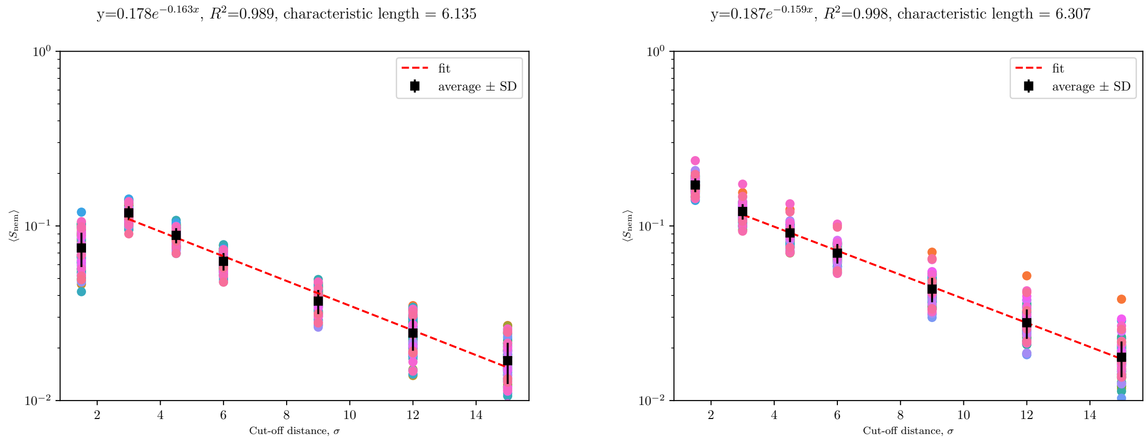


Figure S20: $\langle S_{\mathrm{nem}} \rangle$ due to interstem interactions vs cut-off distance for the last simulation frame for ensembles of simulations with COMRADES score input and stacking that are unconfined (left) and confined (right).

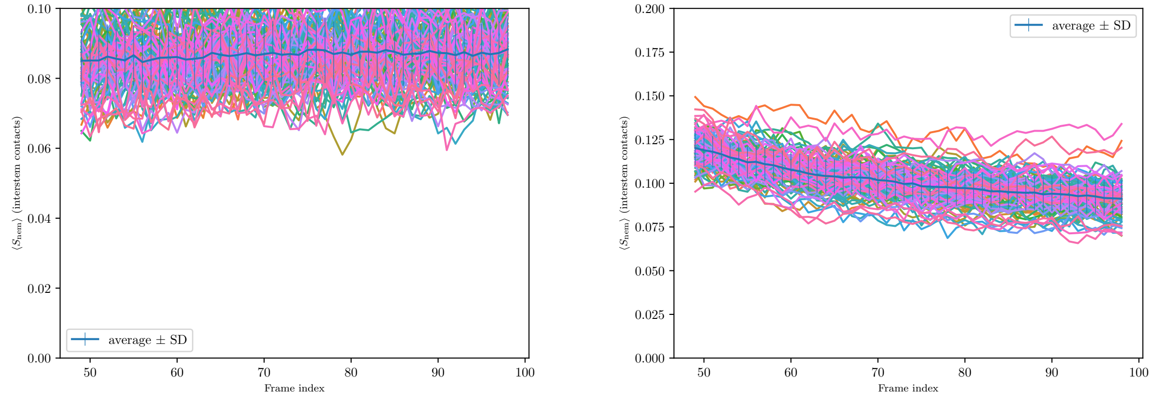


Figure S21: $\langle S_{\mathrm{nem}}(t) \rangle$ arising from interstem interactions for $l_{\mathrm{cut-off}} = 4.5\sigma$ and ensembles of simulations with COMRADES score input and stacking, with unconfined conditions on the left and confined - on the right.

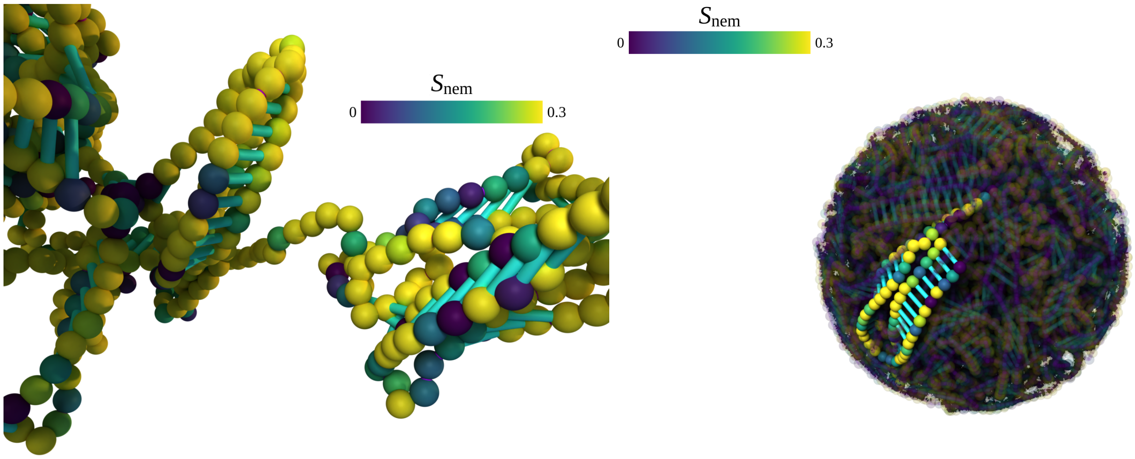


Figure S22: Additional snapshots from the simulations in Figure 4(g,h), colored by the nematic order parameter (cut-off distance $4.5\sigma$). Left: frame 93 of an unconfined simulation; right: frame 98 of a confined simulation.

| Dataset | $l_{\text{nem}}, \sigma$ |
|---|---|
| no confinement, no base-pairing | 2.420 |
| unconfined 3D COMRADES, no stacking | 1.589 |
| unconfined 3D COMRADES with stacking | 2.993 |
| confinement, no base-pairing | 1.465 |
| confined 3D COMRADES, no stacking | 1.546 |
| confined 3D COMRADES with stacking | 2.219 |

Table S5: Characteristic length of the nematic domains $l_{\text{nem}}$ determined from a fit of $\langle S_{\text{nem}} \rangle$ vs. cut-off distance for frame 99 in various types of simulations, see Fig. 4 and Suppl. Fig. S18.

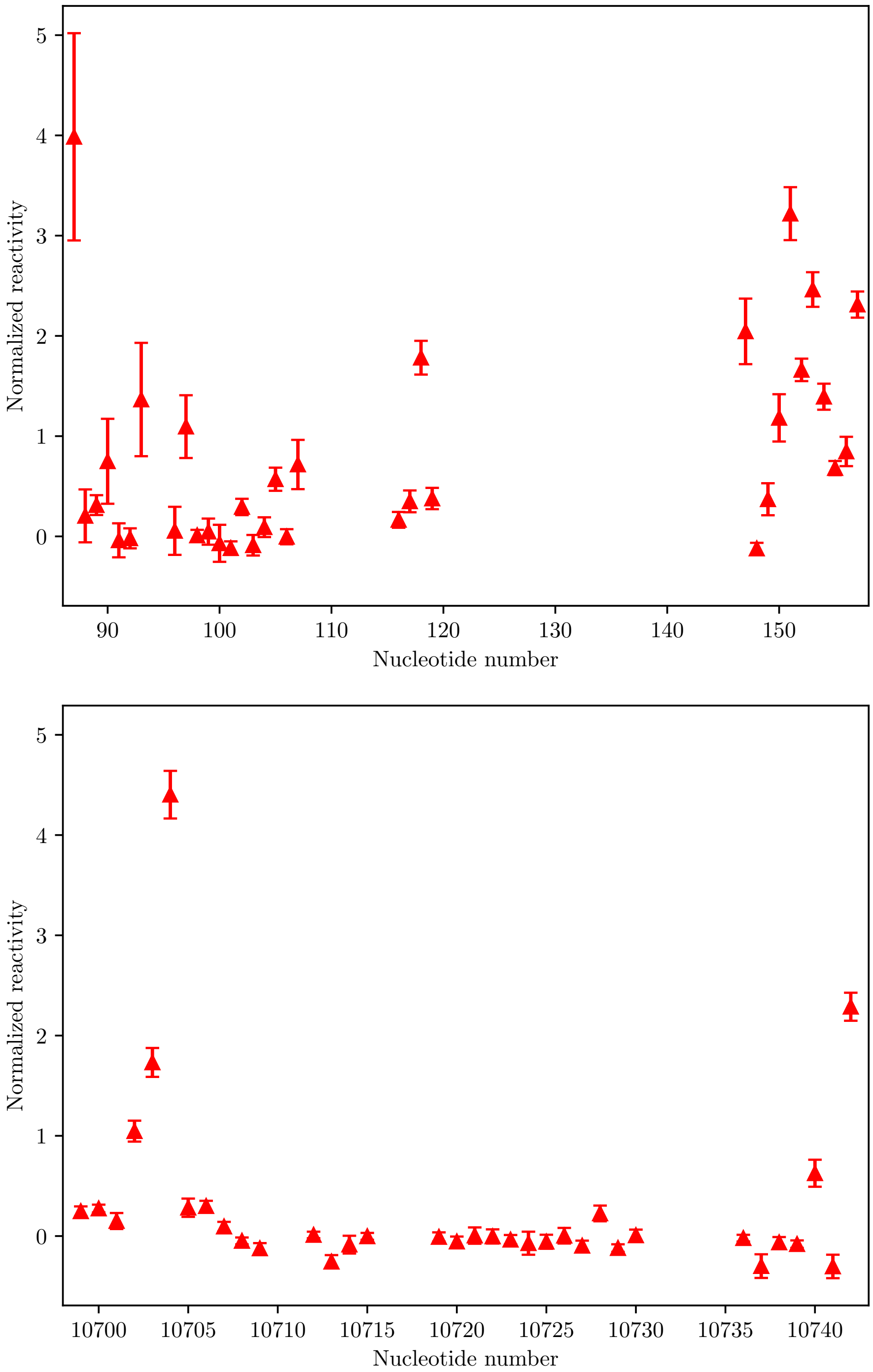


Figure S23: Normalized NAI-MaP reactivity scores obtained from Zika virions (strain KJ776791.2, French Polynesia) by Huber et al. [8] indicates that some of the bases involved in the circularization interaction in Zika have a high reactivity (normalized reactivity > 0.5 [8]) even in the virion, demonstrating that the circularization interaction coexists with other secondary structures. Errorbars: normalized standard error of the reactivity.

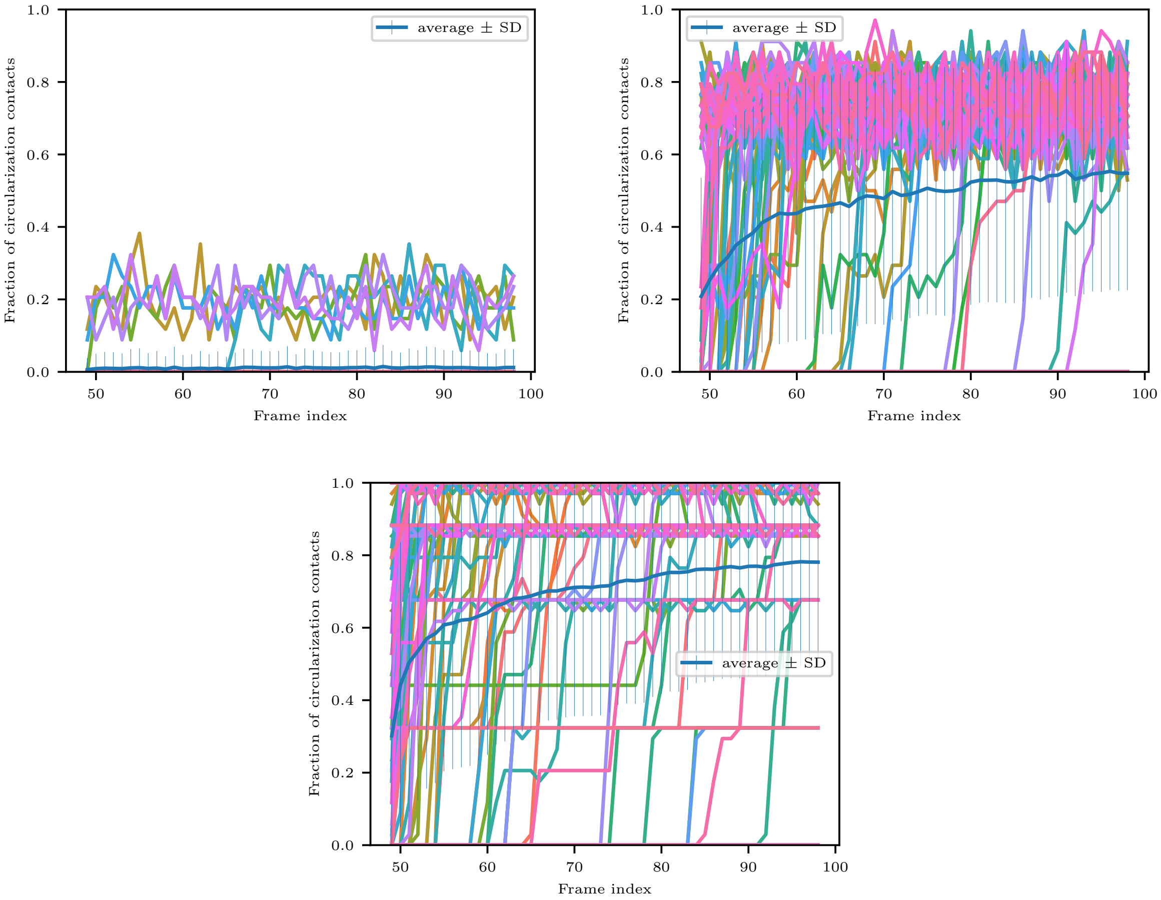


Figure S24: Fraction of circularization contacts vs. time as calculated for ensembles of unconfined simulations (left) and confined ones (right) with no stacking (top row) and with stacking (bottom row, showing data from all runs and complementing Figure 4 from the main text). Ensembles of 100 simulations for all cases.

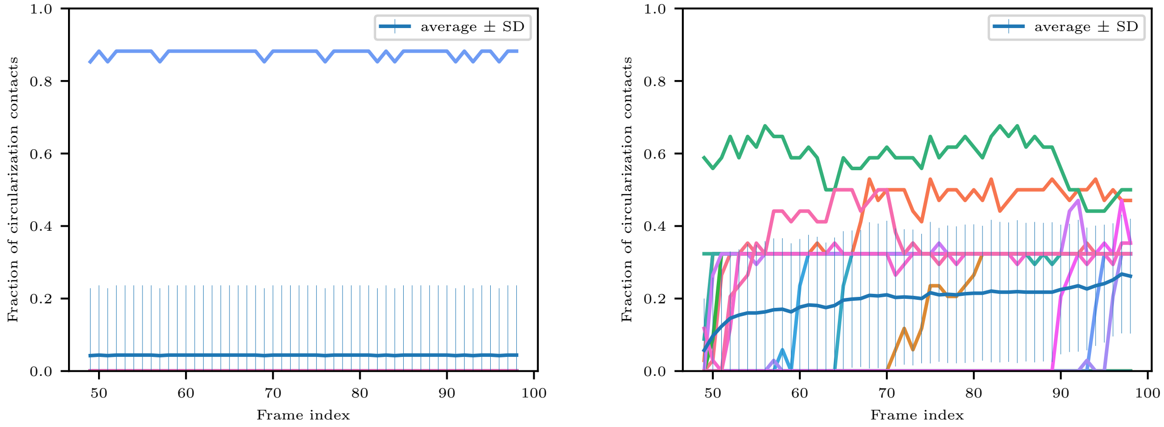


Figure S25: Fraction of circularization contacts vs. time as calculated for ensembles of unconfined simulations using COMRADES scores calculated from experiments with virions as input (left) and unconfined ones with COMRADES score input from in vivo experiments (right). Confinement is the main factor controlling the frequency of circularization. Circularization interactions are rarely seen in unconfined simulations with COMRADES score input based on the data for virions (left), although circularization contacts are common in that dataset (see Figure 4 in the main text and Figure S24). In contrast, circularization contacts are common in confined simulations with in vivo COMRADES score input (frequency of $\sim 25\%$ at long times, right), even those these interactions are comparatively rare in the experimental data. Comparison with the curve for confined simulations with COMRADES score input from experiments with virions shows that circularization is most common in the latter, in which an average of $\sim 75\%$ of circularization contacts are seen at long times (Figure 4). Ensembles of 20 simulations for all cases.

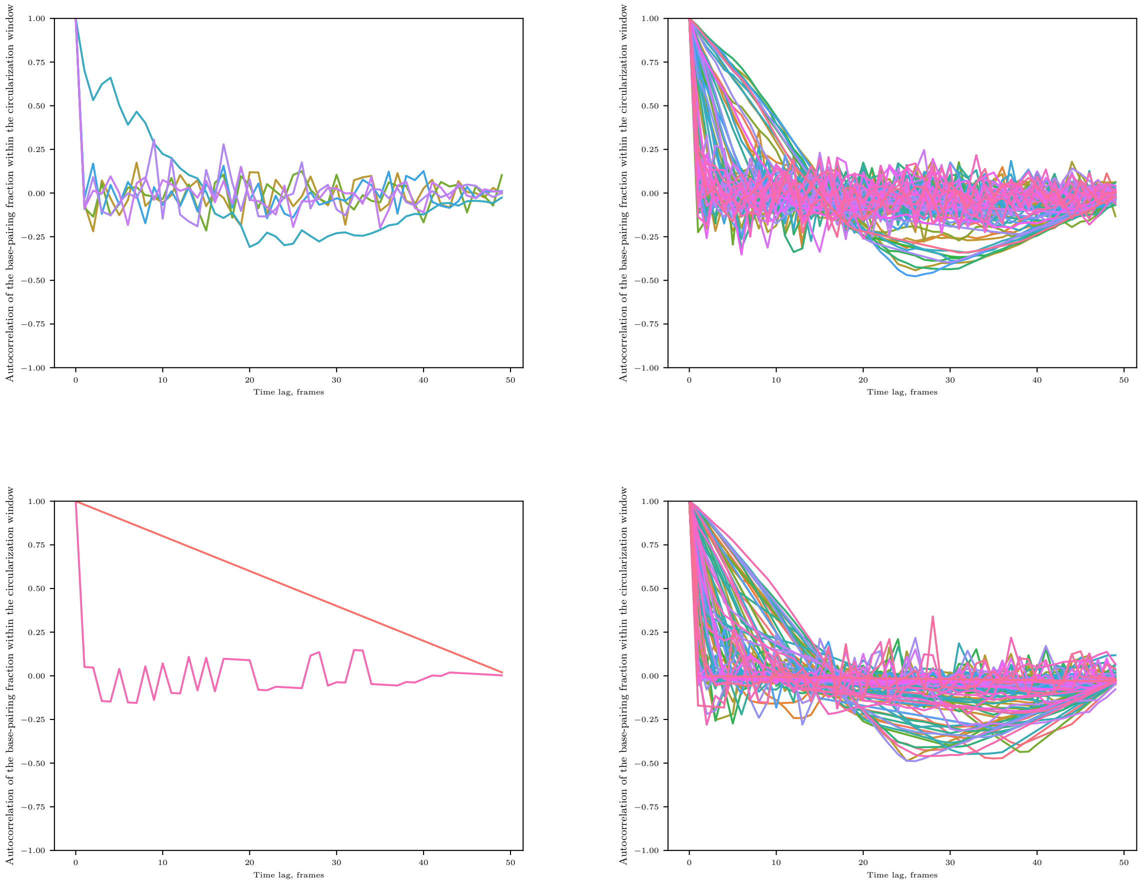


Figure S26: Autocorrelation function of the fraction of circularization contacts vs. time lag as calculated for ensembles of unconfined simulations (left) and confined ones (right), with no stacking in the top row and with stacking in the bottom row. As the circularization fraction is zero for all studied frames in most runs with no confinement, the autocorrelation function is undefined for those runs. Ensembles of 100 simulations for all cases.

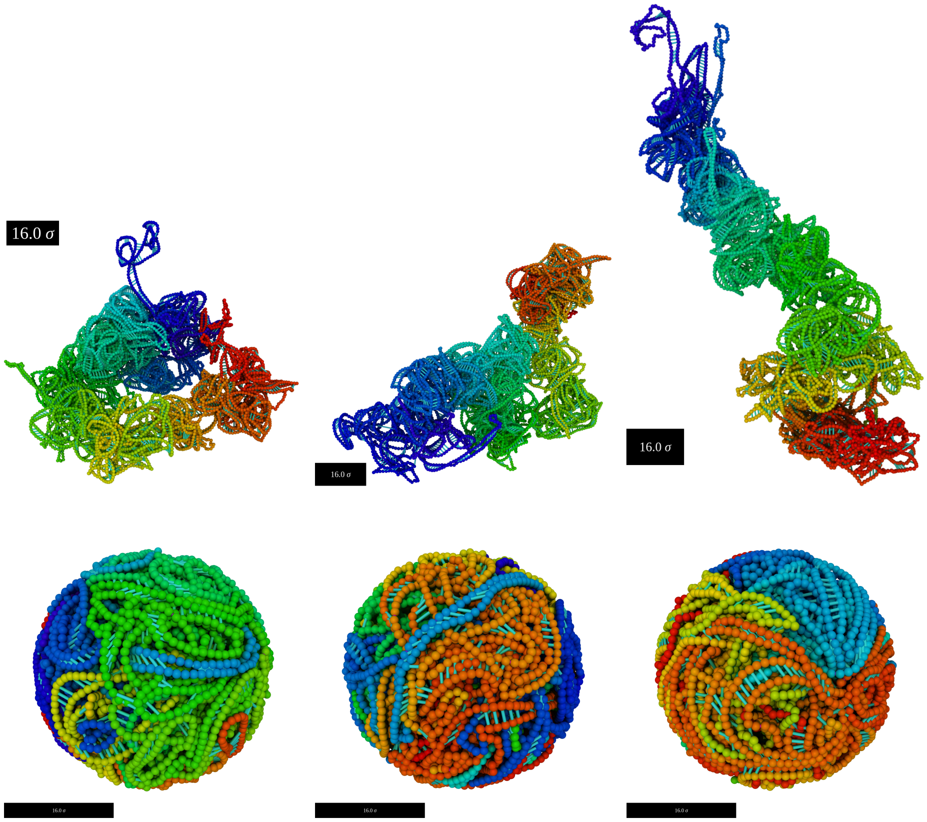


Figure S27: Diversity of the ensemble of 3D conformations in our coarse-grained simulations. Contact patterns, notably the circularization contact, differ between different randomly instantiated runs in the simulation ensemble. Top row: unconfined simulations, bottom row: confined simulations. Scale bar length in simulation snapshots approximately equal to the radius of the inner capsid of the Zika virus.

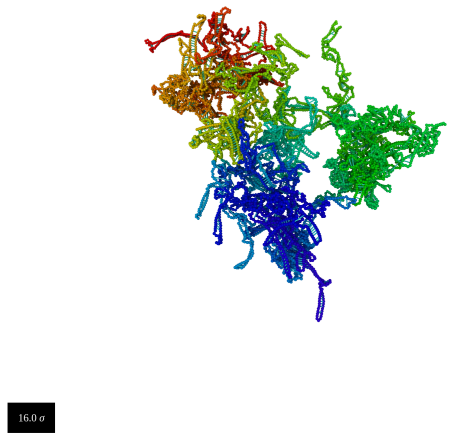


Figure S28: This snapshot from a 3D Vienna simulation displays a more expanded structure than those of 3D COMRADES due to the smaller number of contacts. 3D Vienna predictions exhibit tree-like branched structures that resemble RNAfold-predicted secondary structures expanded into 3D. Scale bar length approximately equal to the radius of the inner capsid of the Zika virus.
